# Interface integrity in septin protofilaments is maintained by an arginine residue conserved from yeast to man

**DOI:** 10.1101/2025.01.29.635471

**Authors:** Benjamin Grupp, Jano Benito Graser, Julia Seifermann, Stefan Gerhardt, Justin A. Lemkul, Jan Felix Gehrke, Nils Johnsson, Thomas Gronemeyer

## Abstract

The septins are conserved, filament-forming, guanine nucleotide binding cytoskeletal proteins. They assemble into palindromic protofilaments which polymerize further into higher-ordered structures that participate in essential intracellular processes such as cytokinesis or polarity establishment. Septins belong structurally to the P-Loop NTPases but, unlike their relatives Ras or Rho, do not mediate signals to effectors through GTP binding and hydrolysis. Biochemical approaches addressing how and why septins utilize nucleotides are hampered by the lack of nucleotide free complexes. Using molecular dynamics simulations, we determined structural alterations and inter-subunit binding free energies in human and yeast septin dimer structures and in their *in silico* generated apo forms.

An interchain saltbridge network in the septin unique β-meander, conserved across all kingdoms of septin containing species, is destabilized upon nucleotide removal, concomitant with disruption of the entire G-interface.

Within this network, we identified a conserved arginine residue, Arg(βb), as the central interaction hub.

The essential role of Arg(βb) for interface integrity was confirmed species-overlapping in human and yeast septins by *in vitro* and *in vivo* experimentation.

## Introduction

Septins are a component of the cytoskeleton (Mostowy and Cossart, 2012). Individual septin subunits form hexa- or octameric linear protofilaments termed rods, serving as building blocks for higher ordered structures such as filaments, gauzes or rings (Bertin *et al*., 2008; Garcia *et al*., 2011; Schneider *et al*., 2013; Kaplan *et al*., 2015). These assemblies are required for essential intracellular processes such as cytokinesis, polarity establishment, or cellular adhesion (Woods and Gladfelter, 2021).

Septins belong to the GTPase superclass of the P-loop NTPases (Leipe *et al*., 2002). The structural topology of the septin GTPase (or short G-) domain resembles a modified Rossmann fold containing all features for GTP hydrolysis such as the P-loop and the switch 1 and switch 2 motifs (Vetter and Wittinghofer, 2001; Cavini *et al*., 2021; Grupp and Gronemeyer, 2022). However, some septins do not hydrolyze their bound nucleotide, classifying them as pseudo-GTPases (Stiegler and Boggon, 2020; Hussain *et al*., 2023). The G-domain is decorated with N- and C-terminal extensions of various length, ranging from a few (C-terminal extension of Cdc10 from yeast) to several hundred amino acids (N-terminal extension of human SEPT9). Another septin-specific structural element of the G-domain is the septin unique element (SUE), consisting of a three-stranded β-meander (SUE-βββ) (Sirajuddin *et al*., 2007; Cavini *et al*., 2021). Septin monomers assemble into apolar, palindromic protofilaments by alternating interactions via their N- and C-termini (NC-interface) and via adjacent G-domains (G-interface), respectively (Sirajuddin *et al*., 2007; Bertin *et al*., 2008; Mendonça *et al*., 2021; Grupp *et al*., 2024), with the SUE-βββ being a central element in G-interface formation.

Phylogenetic analyses grouped animal, fungi and protist septins in 5 groups (Pan *et al*., 2007; Auxier *et al*., 2019; Shuman and Momany, 2022) and recently, two more groups were assigned for septins from algae and one additional for ciliates (Delic *et al*., 2024). In all eight groups conserved residues in the GTPase specific motifs and the SUE can be found whereas residues in the NC interfaces are conserved in groups 1-6a but poorly conserved in the newly assigned groups 6b-8 (Delic *et al*., 2024). This finding led to the assumption that nucleotide binding and hydrolysis is an evolutionary old feature of the septins. What is the role of nucleotide interaction and hydrolysis in septins? Unlike their relatives, the small GTPases from the Ras family, septins do not mediate signals by binding to effector molecules in their active, GTP bound state (Wittinghofer and Vetter, 2011). Experimental data suggest that nucleotide binding and hydrolysis is required for protofilament formation rather than for signal transduction (Sirajuddin *et al*., 2009; Zent *et al*., 2011; Weems *et al*., 2014; Zent and Wittinghofer, 2014; Weems and McMurray, 2017). However, structures of monomeric septins are not yet available (except the yeast septin Cdc11 (Brausemann *et al*., 2016)) and all available dimeric or multimeric septin structures are nucleotide bound. These limitations hampered mechanistical studies aiming at solving the long-standing question how septins employ nucleotides in the process of protofilament formation although such a role was postulated already more than a decade ago (Sirajuddin *et al*., 2009).

Biochemical processes inaccessible to experimentation can sometimes be approached by *in silico* methodology. If sufficient structural information about the protein of interest is available, molecular dynamics (MD) simulations (Zheng *et al*., 2018) can be employed to study for example substrate translocations through channel proteins (Göddeke and Schäfer, 2020), conformational dynamics in small GTPases (Morris *et al*., 2016) or stability of protein complexes (Lemkul and Bevan, 2010). Previously, we employed MD simulations to determine the inter-subunit affinity in human septin G-interface dimers, making use of available crystal structures (Grupp *et al*., 2023). In this experimental setup, the dimer structures and *in silico* generated apo structures were first relaxed in unbiased simulations. These equilibrated structures were subsequently subjected to steered simulations combined with umbrella sampling (Kästner, 2011) to extract eventually free binding energies.

By comparing the nucleotide-bound and apo structures after the unbiased simulations, we identified a conserved arginine residue in the SUE-βββ, Arg(βb), which undergoes a major conformational change (“flipping”) upon nucleotide removal in the investigated catalytically active subunits SEPT2 and SEPT7, accompanied by a reduction of the binding affinity between the respective subunits. We suggested that this arginine flipping upon nucleotide removal is a conserved feature and that the nucleotide plays a pivotal role in G-interface stability. Recently, structures of *S. cerevisiae* septin G-interface dimers became available (Marques da Silva *et al*., 2023; Grupp *et al*., 2024), allowing to perform similar MD simulations with septins from yeast. We present here an improved structure of a yeast septin tetramer with an anisotropic resolution of 2.04 Å, providing an excellent starting point for MD simulations. Based on these simulations we provide herein evidence that the Arg(βb) plays indeed a species-overlapping pivotal role in G-interface formation. Our *in vitro* and *in vivo* experiments in human and yeast cells confirm the prediction that mutation of Arg(βb) results in G-interface disruption.

## Results

### A refined structure of a yeast septin tetramer

We have previously presented the first structure of a septin hetero-tetramer comprised of the yeast-septin subunits Cdc10-Cdc3-Cdc12-Shs1 (PDB-ID 8PFH) (Grupp *et al*., 2024).

Since this complex contained GDP in all subunits, we attempted to crystallize the same complex in the transition state conformation. Inorganic MgF_3_ ^-^ (forming spontaneously in a Mg^2+^ containing buffer from supplemented NaF) should occupy the γ-phosphate binding site and a GDP: MgF_3_ ^-^ preparation of a small GTPase should thus mimic the transition state of the GTPase reaction (Jin *et al*., 2017). We crystallized the hetero-tetramer in the presence of Mg^2+^ and NaF. Diffracting crystals belonged to the space group P-1-2_1_-1 and contained the tetramer Cdc10-Cdc3-Cdc12-Shs1 in the asymmetric unit (Fig. 1A). As in our previous crystallization approach, a filament-like structure forming atypic Cdc10-Shs1 interfaces was present in the crystal. The resulting structure did not reveal the desired transition state conformation but contained again GDP in all subunits (Fig. 1B). However, with anisotropic diffraction limits of 2.017 Å, 3.018 Å and 2.645 Å as determined by STARANISO (Vonrhein *et al*., 2018) (Suppl. Table 1) the overall resolution was significantly improved compared to 8PFH. The structure was deposited in the PDB under identifier 9GD4.

**Figure 1.**
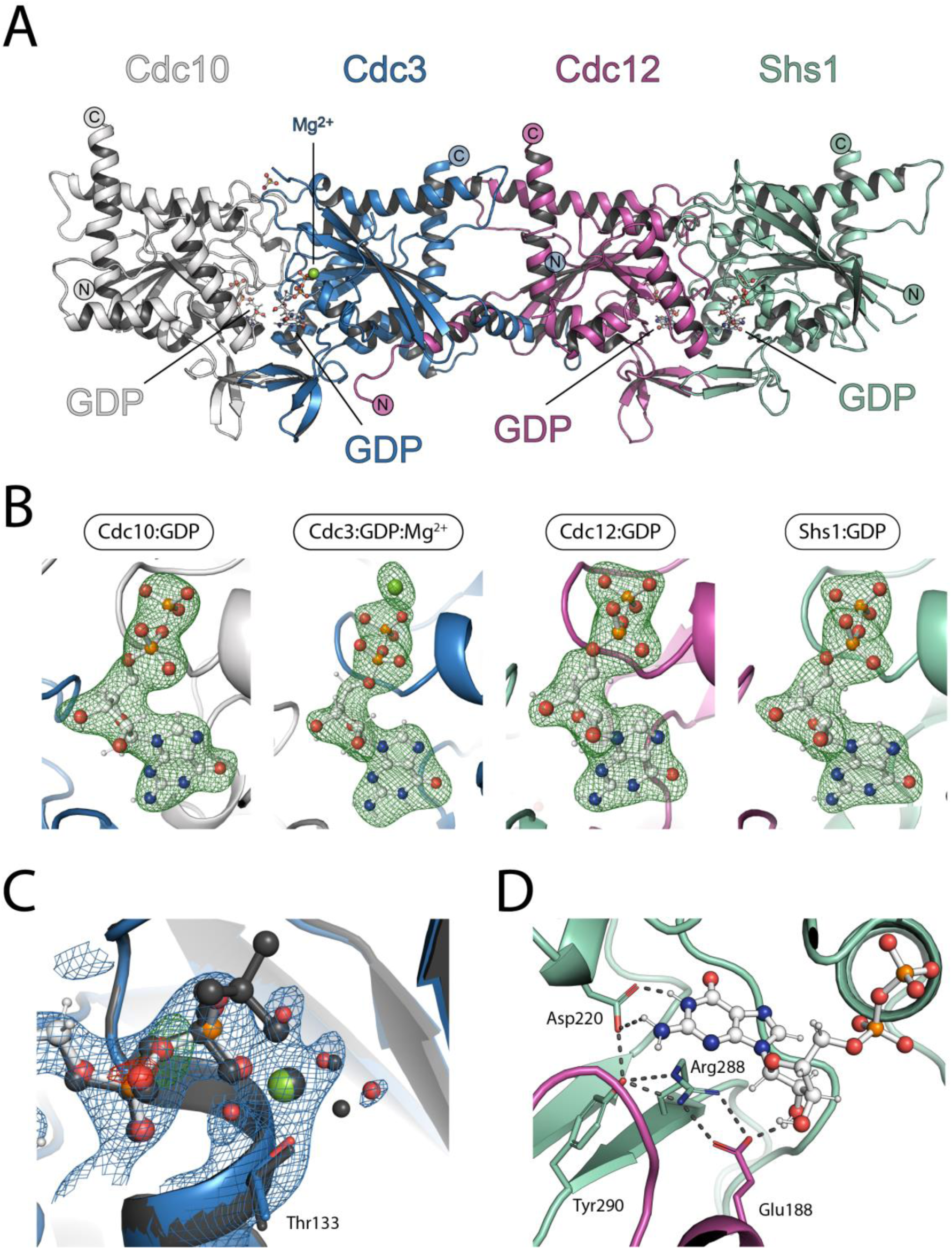
Crystal structure of the tetrameric septin complex. **(A)** The yeast septin complex consisting of Cdc10, Cdc3, Cdc12 and Shs1 in the asymmetric unit of the crystal. N- and C-termini of the subunits are indicated. **(B)** All subunits within the crystal contain GDP as shown by Polder-OMIT maps (contoured at 1.5 σ). **(C)** Nucleotide binding pocket of Cdc3 with a coordinated Mg^2+^ ion. A 2m-DFc electron density map (contoured at 1.5 σ (blue)) and a mFo-DFc electron density map (contoured at +3 σ (green) and -3 σ (red)) is shown. Both maps do not support to build GTP in this subunit. The model was aligned with the Cdc10:GDP-Cdc3:GTP:Mg^2+^ dimer (black) (PDB-ID: 8FWP). **(D)** The conserved hydrogen bond network coordinating the guanine ring of the nucleotide in Shs1. Arg(βb) (here Arg288) forms a triangular hydrogen bond network with the conserved residues Tyr(βb) (Tyr290), Asp(G4) (Asp220) and Glu(α4) (Glu188) from the neighboring subunit Cdc12. Nucleotides are displayed in ball and stick presentation throughout.

The overall fold of this and our previous structure is consequently very similar, with a Cα-RMSD of 0.975 Å. In the nucleotide binding pocket of Cdc3 a coordinated Mg^2+^ ion could be resolved in the same position as the published Cdc3-Cdc10 dimer (PDB-ID: 8FWP) and the Cdc3-Cdc10-Cdc10-Cdc3 tetramer (PDB-ID: 8SGD) (Marques da Silva *et al*., 2023) (Fig 1C.).

### MD simulations

We investigated human septin G-interfaces using molecular dynamics (MD) simulations in previous work and determined the inter-subunit affinities in dependence of the respective nucleotide (Grupp *et al*., 2023). We have identified a highly conserved Arg residue in the septin unique element, Arg(βb), as an essential element for G-interface integrity. Arg(βb) formed a triangular hydrogen bond network with the conserved residues Tyr(βb) and Asp(G4), which altogether coordinated the guanine ring of the nucleotide, and with a Glu(α4) residue from the neighboring subunit. This network is also present in yeast septins (Fig. 1D). In catalytically active human SEPT7 and SEPT2, Arg(βb) flipped by 90° upon nucleotide removal, thereby disrupting the entire hydrogen bond network, suggesting that Arg(βb) plays a pivotal role in G-interface integrity (Grupp *et al*., 2023). Whether this feature is unique for human septins or a common mechanism across species could until now not be answered due to the lack of structural information of septin G-interfaces of other species.

The refined structure of a yeast septin presented here provided us with a crystal structure containing the yeast septin G-interface dimers with a favorable resolution to perform MD simulations, namely Cdc10:GDP-Cdc3:Mg^2+^/GDP and Cdc12:GDP-Shs1:GDP. We generated energy minimized coordinate files for these dimers in their nucleotide-bound conformation and obtained the apo state (Cdc10:apo-Cdc3:apo, Cdc12:apo-Shs1:apo) by *in silico* deletion of the nucleotide. Additionally, we created both dimer sets (with and without nucleotide) with Arg(βb) mutated to alanine. The overall setup of the *in silico* investigation followed the concept of our previous work (Grupp *et al*., 2023). Briefly, each coordinate file structure was temperature- and pressure equilibrated with restraints applied. These input structures were subjected to five independent unbiased simulations without restraints each with a duration of 700 ns, followed by backbone RMSD-based clustering (Daura *et al*., 1998). The central structure in the largest cluster for each system was first examined with respect to structural alterations. Structural integrity of the septin dimers maintained after the unbiased simulations with the Cα root-mean-square deviation (Cα-RMSD) of the relaxed structures remaining below 0.6 nm compared to the energy minimized coordinate structures, with the largest deviations located as expected in the flexible switch loops and loops in the free NC-interfaces (Suppl. Fig. S1).

Structural rearrangements within the Arg(βb) hydrogen bond network after unbiased simulations were quantified by considering distances between involved key residues. The pairwise distances between Arg(βb) and Asp(G4) as well as Arg(βb) and Glu(α4) were extracted from the unbiased simuations of wildtype apo and holo systems and probability densities were obtained by two-dimensional kernel estimation. From the obtained densities relative free energy differences (ΔG) were calculated and plotted as free energy landscapes (Fig. 2). The free energy landscape plot of the apo bound state can be compared with the plot of the respective GDP bound state to illustrate the structural changes in the Arg(βb) hydrogen bond network.

**Figure 2.**
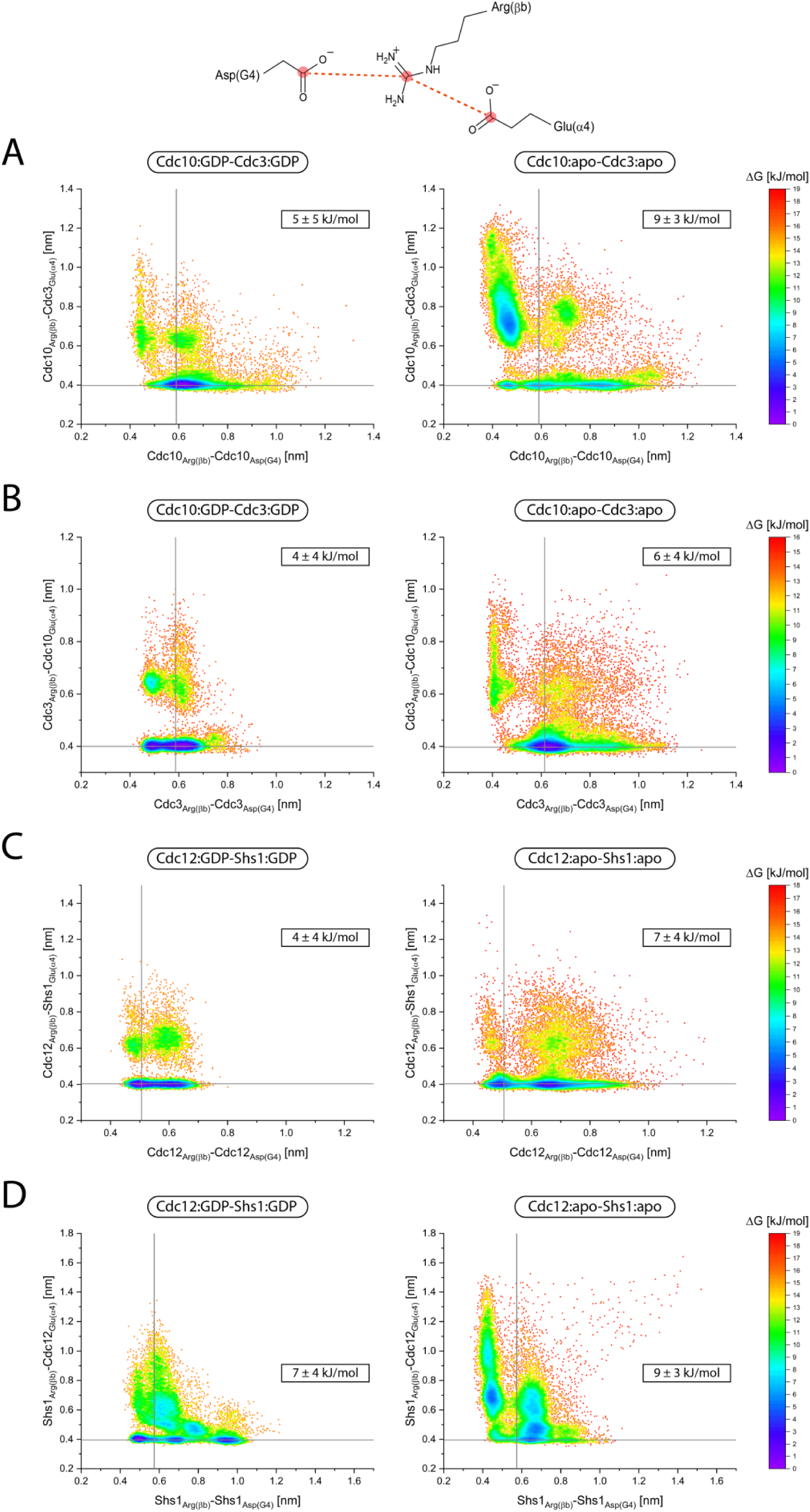
Distances between key residues of the Arg(βb) hydrogen bond network, plotted as free energy landscapes. Pairwise distances from the unbiased simulations for each system were combined and probability densities were calculated by two-dimensional kernel estimation, from which relative free energy differences (ΔG) were calculated. The state with the highest calculated density across corresponding apo and holo systems served as a reference point. ΔG relative to the global energy minimum for both systems is shown as a color gradient. For each system, the average ΔG and its standard deviation is given in the inserts. Reference lines in the plots indicate the distances present in the initial energy minimized coordinate structures, the intersection represents the observed distance pair. Each dot represents one distance pair. Free energy landscape plots are shown for distances between **(A)** Cdc10Arg(βb) - Cdc10Asp(G4) (x-axis) and Cdc10Arg(βb) - Cdc3Glu(α4) (y-axis), **(B)** Cdc3Arg(βb) - Cdc3Asp(G4) (x-axis) and Cdc3Arg(βb) - Cdc10Glu(α4), **(C)** Cdc12Arg(βb) - Cdc12Asp(G4) (x-axis) and Cdc12Arg(βb) - Shs1Glu(α4) (y-axis), and **(D)** Shs1Arg(βb) - Shs1Asp(G4) (x-axis) and Shs1Arg(βb) - Cdc12Glu(α4) (y-axis). For all panels, the GDP-bound state is shown on the left and the apo state is shown on the right.

Upon removal of the nucleotides from the Cdc10-Cdc3 dimer, the hydrogen bond networks became disturbed, as shown by a pairwise increased scattering of both distances between the respective residues of the hydrogen bond network. The average ΔG for the Cdc10Arg(βb) apo and Cdc3Arg(βb) apo system was increased compared to the respective GDP system, which confirms an overall destabilization of the interface in the apo state (Fig. 2A, B).

For Cdc10Arg(βb), the global energy minimum in the apo bound state was shifted by approx. 0.1 nm towards a smaller Arg(βb)-Asp(G4) spacing but 0.3 nm towards a higher Arg(βb)-Glu(α4) spacing, which suggests a rapid destabilization of the initially observed network over the time course of the simulation. An increased scattering along both axes could be seen, indicative of a weakening of both the Arg(βb)-Glu(α4) and Arg(βb)-Asp(G4) interactions (Fig. 2A).

For Cdc3Arg(βb), the local energy minimum was not displaced in the apo state, suggesting a longer dwell time in the initial state of the simulation. As for Cdc10, both the Arg(βb)-Glu(α4) and Arg(βb)-Asp(G4) interactions weakened, as indicated by the increased number of low energy state observations with higher distances (Fig. 2B).

Removal of the nucleotide from the Cdc12-Shs1 dimer resulted also in a global destabilization of the interface, indicated by a pairwise increased scattering of both distances and an increase of the average ΔG of both apo systems (Fig. 2C, D).

The Arg(βb)-Asp(G4) interaction, but not the Arg(βb)-Glu(α4) interaction was weakened for Cdc12Arg(βb) (Fig. 2C). On the contrary, this weakening is more prominent for the Arg(βb)-Asp(G4) interaction than the Arg(βb)-Glu(α4) interaction, as determined by the appearance of a second energy minimum.

To complement the free energy landscape approach with a more global view on structural deformations of the whole SUE-βββ, we performed a backbone RMSD analysis. We compared the RMSD distribution of the SUE-βββ, obtained from the pooled unbiased simulations of each septin subunit, with the respect to the equilibrated input structure (Suppl. Fig. S2). The 75 % quantile and the maximum RMSD value was shifted for any introduced modification (removal of the nucleotide, Arg(βb) mutation or a combination of both) in any sepin subunit. This behavior indicated an increased probability of a structural deformation in the SUE-βββ. Compared to the other subunits, Shs1 displayed the most uniform violin plot shapes and smallest shift of the 75 % quantile, which suggests that the SUE-βββ of this subunit is less prone to structural alterations.

The structure of the central clustered frame for each system was subsequently employed for the determination of inter-subunit affinities. We describe in the following in more detail the structural alterations of these structures compared to the respective coordinate file structures.

In the unmodified Cdc10-Cdc3 and Cdc12-Shs1 dimers, the hydrogen bond network around Arg(βb) remained intact (Suppl. Fig S3A and S4A). Upon removal of the nucleotides from the Cdc10-Cdc3 dimer, flipping of the Arg(βb) could be seen in both subunits (Suppl. Fig. S3B). Additionally, the hydrogen bonds connecting the βb and βc strands in the Cdc10-SUE became distorted. As a result, Cdc10-SUE-βββ partially dissociated from the Cdc3-SUE, indicated by altered positioning of Trp(βb) relative to Cdc10 (Suppl. Fig. S3B). Introduction of the Arg(βb)-Ala mutation in both subunits in the nucleotide-bound state left the Asp(G4) hydrogen bonding to the guanine ring intact. Otherwise, as in the apo form, the hydrogen bonds connecting the βb and βc strands in the Cdc10-SUE became perturbed, resulting in partial dissociation of the Cdc10-SUE-βββ from the Cdc3-SUE (Suppl. Fig S3C.).

Removal of the nucleotide from the Cdc12-Shs1 dimer resulted only in a partial disruption of the hydrogen bond network, corresponding to the slight deformation of Arg(βb), but without flipping (Suppl. Fig. S4B). Larger conformational displacements within the SUE occurred, as indicated by the partial dissociation of the Cdc12-SUE-βββ from Shs1. Furthermore, the two Tyr(βb) residues of both subunits tilted towards each other (Suppl. Fig. 4B). Introduction of the Arg(βb)-Ala mutation lead additionally to disruption of the contact between the conserved Trp(βb) residues of both subunits (Suppl. Fig S4C). For both Cdc10-Cdc3 and Cdc12-Shs1 dimers, we also tested the Arg(βb)-Ala mutation in the apo form. No further structural alterations could be observed (Suppl. Fig. S3D and S4D).

To determine inter-subunit affinities, the central structure in the largest cluster from the unbiased simulations was extracted and protein-protein binding free energy profiles were determined by COM-pulling and umbrella sampling (Lemkul and Bevan, 2010; Patel and Ytreberg, 2018; Grupp *et al*., 2023). Briefly, the filament axis of each structure was aligned with the x-axis, both subunits were separated along the x-axis in COM-pulling simulations and different conformations along the reaction coordinate were extracted. COM separation distances ranged from 3.5 nm (starting conformation) to approximately 8.5 nm.

Binding free energies (ΔG) for the different dimers were calculated by the weighted histogram analysis method (Hub *et al*., 2010). The corresponding free energy profiles are shown in Suppl. Fig. S5A and S5B. Binding free energy in the Cdc12-Shs1 G-interface was increased by approximately 50 kJ mol^-1^ upon nucleotide removal, introduction of the Arg(βb)-Ala mutation, or both (Table1, Suppl. Fig. 5C). In the Cdc10-Cdc3 dimer, ΔG increased by approximately 30 kJ mol^-1^ upon nucleotide removal and by approximately 50 kJ mol^-1^ upon combined introduction of the Arg(βb)-Ala mutation and nucleotide removal (Table1, Suppl. Fig. 5C). Introduction of the mutation alone did not result in an increase of ΔG. Consequently, these results suggest that nucleotide removal leads to a destabilization of the yeast septin interfaces, reflected in the increased ΔG, in agreement with our previous observations for human septins (Grupp *et al*., 2023).

In the following we present our efforts to investigate the role of Arg(βb) *in vivo* and *in vitro* both in human and yeast septins.

**Table 1.** Binding free energies for the Cdc10-Cdc3 and Cdc12-Shs1 systems.

| System | $\Delta G$ [kJ*mol<sup>-1</sup>] | $\Delta\Delta G$ [kJ*mol<sup>-1</sup>] |
| --- | --- | --- |
| *Cdc10-Cdc3* |  |  |
| Cdc10:GDP-Cdc3:GDP | -137 ± 11 | - |
| Cdc10:apo-Cdc3:apo | -103 ± 14 | 33 ± 1 |
| Cdc10<sub>R251A</sub>:GDP-Cdc3<sub>R360A</sub>:GDP | -135 ± 21 | 2 ± 24 |
| Cdc10<sub>R251A</sub>:apo-Cdc3<sub>R360A</sub>:apo | -89 ± 17 | 45 ± 20 |
| *Cdc12-Shs1* |  |  |
| Cdc12:GDP-Shs1:GDP | -189 ± 19 | - |
| Cdc12:apo-Shs1:apo | -133 ± 9 | 56 ± 21 |
| Cdc12<sub>R263A</sub>:GDP-Shs1<sub>R288A</sub>:GDP | -138 ± 35 | 51 ± 39 |
| Cdc12<sub>R263A</sub>:apo-Shs1<sub>R288A</sub>:apo | -133 ± 21 | 56 ± 28 |

### Arg(βb) is essential for filament incorporation of SEPT9 and SEPT9 mediated cell migration

We have recently created a SEPT9 knockout (ko) in human dermal fibroblasts (Hecht *et al*., 2024). This cell line provides an excellent tool to study the role of Arg(βb) (Arg516 in SEPT9) in the context of G-interface formation between SEPT7 and SEPT9 inside the living cell.

We generated an EGFP-SEPT9(R516A) construct and additionally EGFP-SEPT9(W520A). Trp520 in SEPT9 is an absolute conserved residue in the SUE that is essential for G-interface integrity. Mutating this Trp to Ala should disrupt the SEPT9-SEPT7 G-interface as shown for SEPT2 (Sirajuddin *et al*., 2007).

Both mutations and wildtype EGFP-SEPT9 were transfected into the ko cell line and stably expressing cell lines were selected. The intracellular localization of the EGFP-SEPT9 constructs was monitiored using fluorescence microscopy with the endogenous septin cytoskeleton stained with an anti-SEPT8 antibody and actin stained with Phalloidin. EGFP-SEPT9 was incorporated readily into endogenous septin fibers, displaying the typical phenotype of wild-type septin filaments including partly colocalization with actin stress fibers as described by us and others (Dolat *et al*., 2014; Spiliotis *et al*., 2016; Hecht *et al*., 2024) (Fig. 3A, upper panel). The Trp520 mutant did only weakly if at all align with the endogenous septin cytoskeleton (Suppl. Fig S6.) We found a similar result for cells expressing EGFP-SEPT9(R516A). Localization of this construct was mostly diffuse or in filament-like formations concentrated at the plasma membrane. Colocalization with the endogenous septin- and actin cytoskeleton was only occasionally observed (Fig. 3A, lower panel). Western blot analysis employing an anti-SEPT9 antibody of extracts from 1306 WT cells, SEPT9 ko cells and cells expressing EGFP-SEPT9 or EGFP-SEPT9(R516A) confirmed lower expression levels in the soluble extract and degradation products for EGFP-SEPT9(R516A) (Suppl. Fig S7). We conclude that the not-incorporated protein becomes either insoluble or unstable, leading to degradation inside the cell or during extraction.

**Figure 3.**
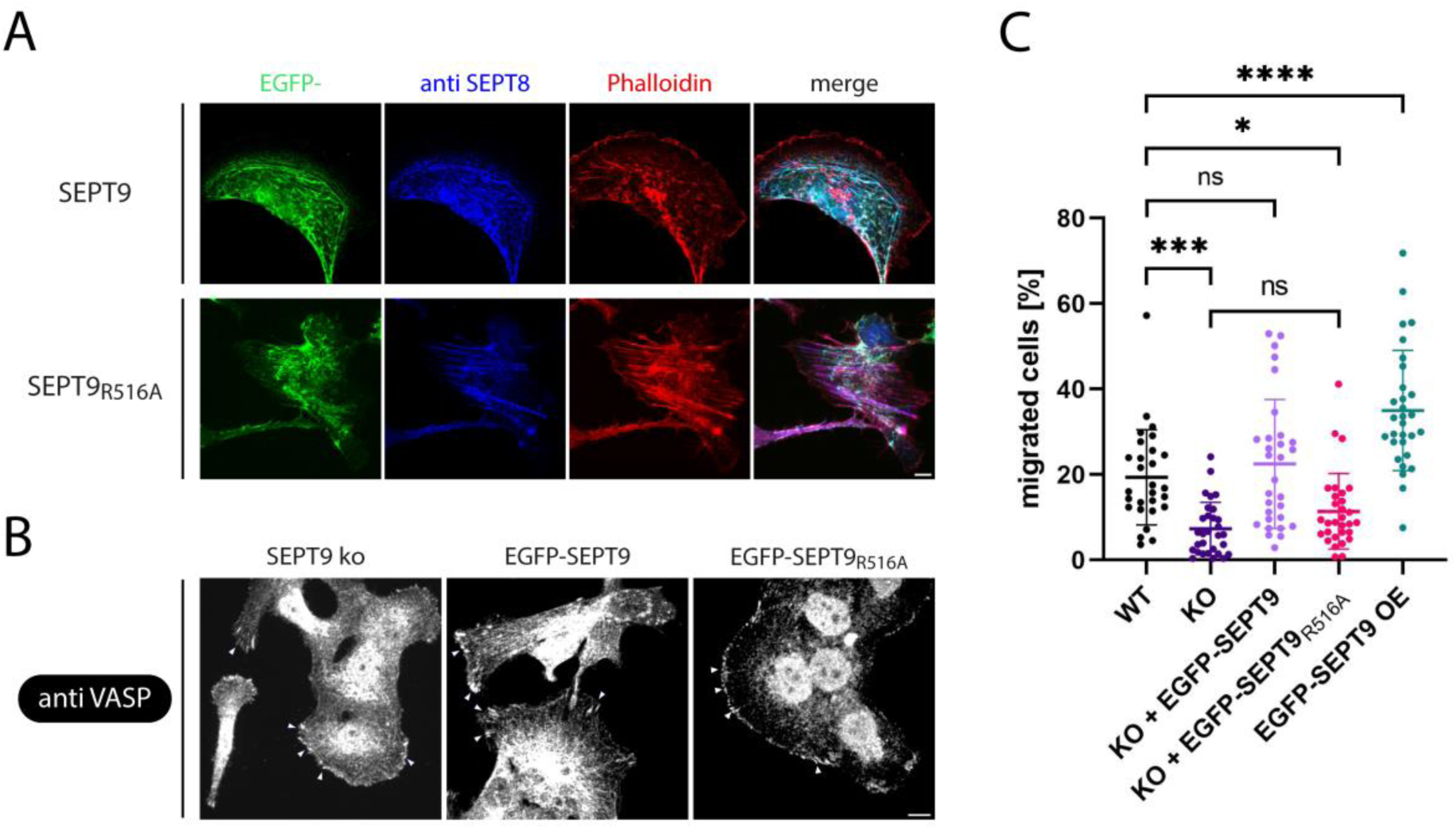
Arg(βb) is essential for filament incorporation of SEPT9 *in vivo* and SEPT9 mediated cell migration. **(A)** Human 1306 fibroblasts with SEPT9 ko, stably overexpressing EGFP-SEPT9 (upper panel) or EGFP-SEPT9(R516A) (lower panel). The endogenous actin was stained with Phalloidin and the endogenous septin cytoskeleton was visualized with an anti-SEPT8 antibody. Note that the majority of EGFP-SEPT9(R516A) is not incorporated into endogenous septin structures. In most observed cells no colocalization at all could be detected. Shown are sum intensities projections. Scale bar 10 µM. **(B)** 1306 fibroblasts with SEPT9 ko exhibit restricted filopodia as indicated by staining with an anti-VASP antibody (right). Stable overexpression of EGFP-SEPT9 in these cells restores filipodia length (middle) whereas EGFP-SEPT9(R516A) overexpressing cells maintain the ko phenotype (right). Filopodia are highlighted with white arrowheads. Shown are max intensity projections. Scale bar 10 µM. **(C)** Boyden chamber assay to monitor cellular migratory behavior. Overexpression of EGFP-SEPT9 in SEPT9 ko cells restores migration rate to wildtype level whereas EGFP-SEPT9(R516A) overexpressing cells maintain the migration rate of ko cells. Significance values were caculated by one-way ANOVA followed by Tukey post hoc test with N=28 to N=30 collected from three independent experiments. Data are depicted as scattered dot plot with means ± SD.

SEPT9 is functionally linked to focal adhesion maturation and cellular migration (Fuechtbauer *et al*., 2011; Dolat *et al*., 2014; Lam and Calvo, 2019). Accordingly, SEPT9 ko fibroblast cell line exhibited restricted, shortened filopodia (Hecht *et al*., 2024) (Fig. 3B, left). Expression of EGFP-SEPT9 in SEPT9 ko cells restoreed filipodia length (Fig. 3B, middle) whereas EGFP-SEPT9(R516A) overexpressing cells exhibited a similar, restricted filipodia shape as the ko cells (Fig. 3B, right).

In line with these findings, SEPT9 ko cells barely migrated in a transwell-assay whereas overexpression of SEPT9 in WT cells increased the migration rate (Hecht *et al*., 2024) (Fig. 3C). Stable overexpression of EGFP-SEPT9 in SEPT9 ko cells restored the migration rate to wild type level whereas cells stably overexpressing EGFP-SEPT9(R516A) displayed low migration rates close to those of the ko cells (Fig 3C).

### Arg(βb) is essential for yeast septin function in vivo

We investigated next the impact of Arg(βb) on subunit association in yeast septins. To test if Arg(βb) is necessary for protofilament formation *in vivo*, we generated yeast strains each with a knock-out for one mitotic septin (Cdc10, Cdc3, Cdc12, Cdc11, Shs1). Each strain expressed its genomically deleted septin from an extrachromosomal rescue plasmid containing an uracil prototrophy. The cells were subsequently transformed with a second plasmid, containing either a wildtype copy of the deleted septin, a copy bearing the Arg(βb)-Ala mutation, or containing no additional coding sequence. Serial dilutions of each strain were spotted onto selective media containing 5-fluoorotic acid (FOA), forcing the cells to drive out the rescue plasmid. The plates were incubated at different temperatures and colony growth was documented. Knock out of each septin except of Shs1 was lethal, indicated by complete growth inhibition of the empty plasmid control on FOA medium (Fig. 4). Arg(βb) to Ala mutation was lethal for cells expressing the corresponding cdc11 and cdc12 alleles at all tested temperatures. The same mutation resulted only in temperature sensitivity (restricted growth at 37 °C) in cdc10 and cdc3 alleles. Strains containing the mutation in these subunits grew normal at 30 °C (Fig. 4).

**Figure 4.**
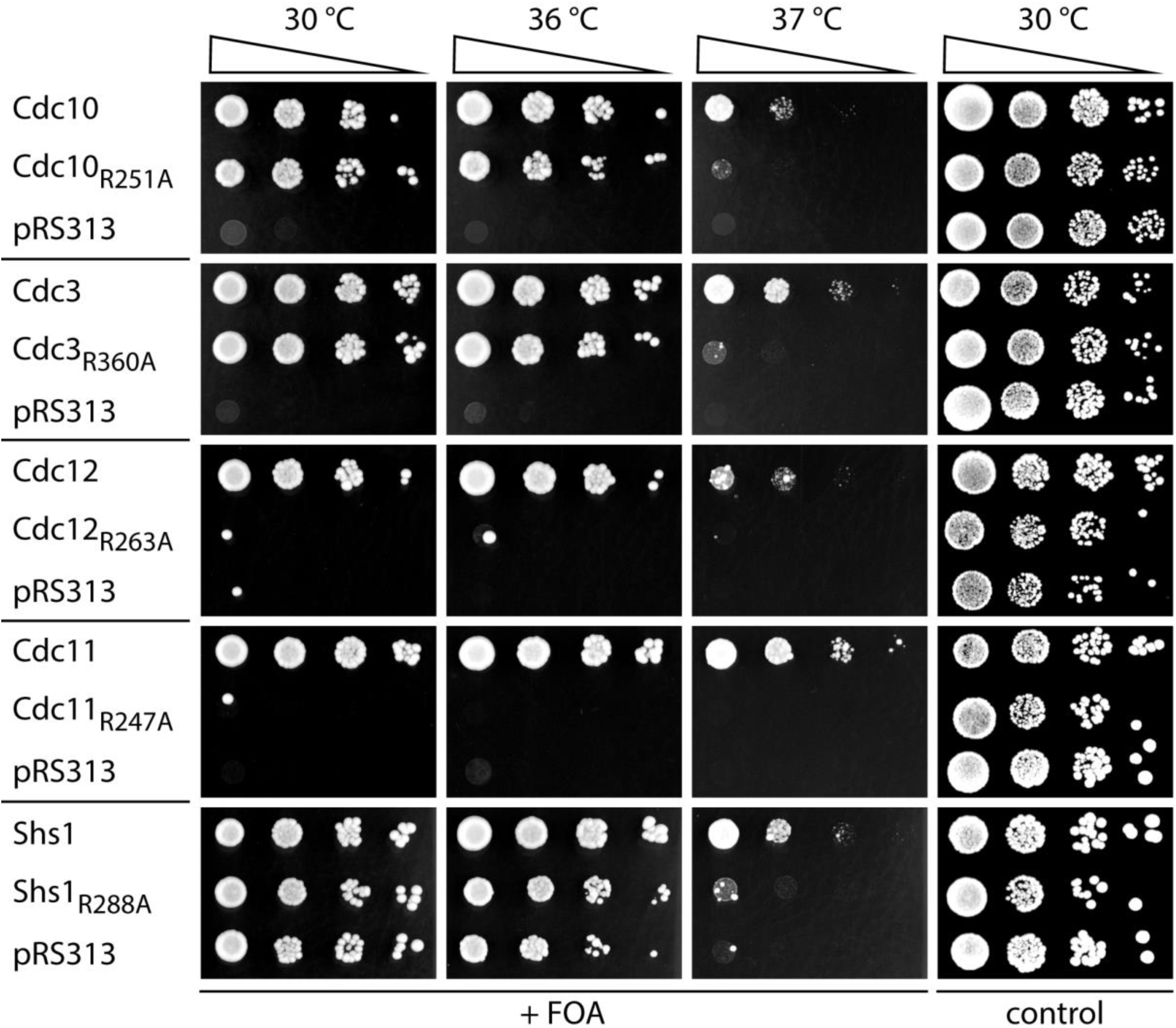
Growth assay with yeast expressing septins with Arg(βb) mutation. Yeast cells with a ko for each one septin (from top to bottom: Cdc10, Cdc3, Cdc12, Cdc11, Shs1) were transformed either with wildtype, Arg(βb)-Ala mutation or empty plasmid. Serial dilutions of each strain were spotted onto selective media containing 5-fluoorotic acid (FOA), forcing the cells to drive out a previously introduced rescue plasmid containing an Ura3 marker. Note that the Arg(βb) mutation results in temperature sensitivity for all tested strains and in lethality even at permissive temperature for Cdc12 and Cdc11.

To validate this finding, we constructed GFP fusion proteins with and without Arg(βb) to Ala mutation of each septin subunit and tested the intracellular localization in two different yeast strain backgrounds (Fig. 5 and Suppl. Fig. S8). The budneck localization was quantified in relation to a Shs1-mCherry marker signal (Fig. 5A). Consistent with the growth assay, Cdc12, Cdc11 and Shs1 containing the Arg(βb) mutation failed to localize at the budneck.

**Figure 5.**
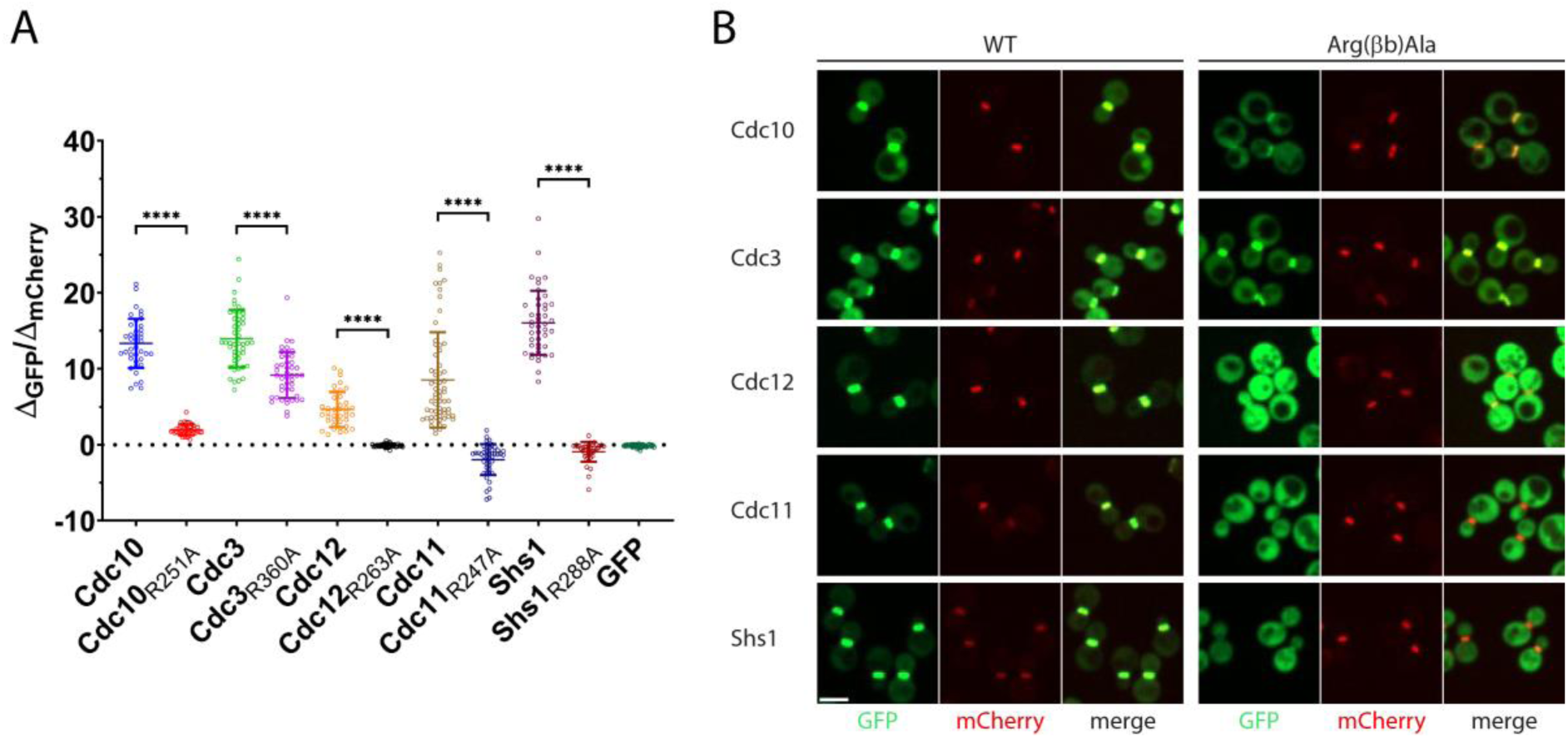
Localization of septins containing Arg(βb) mutation inside the living yeast cell. Septins with and without Arg(βb)-Ala mutation were expressed as GFP fusion protein in yeast cells containing Shs1-mCherry as septin marker. **(A)** Quantification of the bud neck signal (in relation to the marker signal) of the GFP fusion constructs. Only cells in G2/M phase with a clearly detectable Shs1-mCherry signal at the bud neck were considered. Bud neck localization is entirely abolished for Arg(βb) mutation in Cdc12, Cdc11 and Shs1 and reduced for Cdc10 and Cdc3. Significance values were calculated by an unpaired t-test with Welch’s correction with at least N=39 cells. Data are depicted as scattered dot plot with means ± SD. **(B)** Representative microscopy images for each GFP tagged septin subunit as wildtype or Arg(βb)-Ala mutation. In the mCherry channel the Shs1-mCherry marker is shown. Note the diffuse localization of GFP-Cdc12, GFP-Cdc11 and GFP-Shs1 each bearing the Arg(βb)-Ala mutation. Shown are sum intensities projections. Scale bar 5 µM.

The GFP signal was distributed unspecifically in the cytoplasm (Fig. 5B). Budneck localization with significantly reduced fluorescence intensity was still detectable for the Arg(βb) mutants in Cdc10 and especially Cdc3 (Fig. 5A, B).

### Arg(βb) is essential for G-interface formation in septin dimers in vitro

We asked next if our findings could be reproduced *in vitro* using recombinantly expressed G-interface dimers. The central G-interface dimers of human hexameric- or octameric rods, the SEPT7 homodimer and the SEPT7-SEPT9 heterodimer, respectively, were employed for this purpose. The truncated G-domains of SEPT7 and SEPT9, each with or without mutation of the Arg(βb) to Ala, were expressed from two compatible plasmids (Table 2) within the same cell. Since both G-domains have a comparable molecular weight, SEPT7 was fused to MBP, enabling its differentiation from SEPT9 on a SDS-PAGE gel. SEPT9 was fused to a His_6_-tag for purification. For the SEPT7 homodimer, His_6_-SEPT7 constructs with and without Arg(βb) to Ala mutation were expressed.

**Table 2.** List of plasmids used in this study. pAc-MBP, pAc and pES are an in-house constructed, pET15A based plasmids for the expression of MBP- or His_6_ tagged proteins in E. coli. fl -full length, aa – amino acid residue

|  | Construct | Vector backbone | Insert | Source |
| --- | --- | --- | --- | --- |
| #1 | EGFP-SEPT9 | pEGFP-C2 | fl | E. Spiliotis |
| #2 | EGFP-SEPT9(R516A) | pEGFP-C2 | fl | This study |
| #3 | EGFP-SEPT9(W520A) | pEGFP-C2 | fl | This study |
| #4 | DsRed-SEPT9 | pDsRed-Monomer C1 | fl | M. Hecht |
| #5 | EGFP-SEPT2 | pEGFP-C2 | fl | M. Hecht |
| #6 | His<sub>6</sub>-SEPT9G | pES | aa 295-567 | (Hecht *et al.*, 2024) |
| #7 | His<sub>6</sub>-SEPT9G(R516A) | pES | aa 295-567 | This study |
| #8 | MBP-SEPT7G | pAc-MBP | aa 48-318 | This study |
| #9 | MBP-SEPT7G(R265A) | pAc-MBP | aa 48-318 | This study |
| #10 | His<sub>6</sub>-MBP-SEPT7G | pAc | aa 48-318 | This study |
| #11 | His<sub>6</sub>-MBP-SEPT7G(R265A) | pAc | aa 48-318 | This study |
| #12 | His<sub>6</sub>-SEPT7G | pES | aa 48-318 | This study |
| #13 | His<sub>6</sub>-SEPT7G(R265A) | pES | aa 48-319 | This study |
| #14 | His<sub>6</sub>-MBP-Cdc10G | pAc | aa 30-end | This study |
| #15 | His<sub>6</sub>-MBP-Cdc10G(R251A) | pAc | aa 30-end | This study |
| #16 | His<sub>6</sub>-MBP-Cdc12G | pAc | aa 32-314 | This study |
| #17 | His<sub>6</sub>-MBP-Cdc12G(R263A) | pAc | aa 32-314 | This study |
| #18 | His<sub>6</sub>-Shs1G | pET-Duet1 | aa 21-339 | This study |
| #19 | His<sub>6</sub>-Shs1G(R288A) | pET-Duet1 | aa 21-339 | This study |
| #20 | His<sub>6</sub>-Cdc11G | pET-Duet1 | aa 20-298 | This study |
| #21 | His<sub>6</sub>-Cdc11G(R247A) | pET-Duet1 | aa 20-298 | This study |
| #22 | His<sub>6</sub>-Cdc3G | pET-Duet1 | aa 117-410 | This study |
| #23 | His<sub>6</sub>-Cdc3G(R360A) | pET-Duet1 | aa 117-410 | This study |
| #24 | P<sub>MET</sub>-Cdc10 | pRS316 | fl | (Grupp *et al.*, 2024) |
| #25 | P<sub>MET</sub>-Cdc3 | pRS316 | fl | This study |
| #26 | P<sub>MET</sub>-Cdc12 | pRS316 | fl | This study |
| #27 | P<sub>MET</sub>-Cdc11 | pRS316 | fl | This study |
| #28 | P<sub>MET</sub>-Shs1 | pRS316 | fl | This study |
| #29 | P<sub>MET</sub>-Cdc10 | pRS313 | fl | (Grupp *et al.*, 2024) |
| #30 | P<sub>MET</sub>-Cdc10(R251A) | pRS313 | fl | This study |
| #31 | P<sub>MET</sub>-Cdc3 | pRS313 | fl | This study |
| #32 | P<sub>MET</sub>-Cdc3(R360A) | pRS313 | fl | This study |
| #33 | P<sub>MET</sub>-Cdc12 | pRS313 | fl | This study |
| #34 | P<sub>MET</sub>-Cdc12(R263A) | pRS313 | fl | This study |
| #35 | P<sub>MET</sub>-Cdc11 | pRS313 | fl | This study |
| #36 | P<sub>MET</sub>-Cdc11(R247A) | pRS313 | fl | This study |
| #37 | P<sub>MET</sub>-Shs1 | pRS313 | fl | This study |
| #38 | P<sub>MET</sub>-Shs1(R288A) | pRS313 | fl | This study |
| #39 | P<sub>MET</sub>-Cdc10-GFP | pRS306 | fl | This study |
| #40 | P<sub>MET</sub>-Cdc10(R251A)-GFP | pRS306 | fl | This study |
| #41 | P<sub>MET</sub>-GFP-Cdc3 | pRS306 | fl | This study |
| #42 | P<sub>MET</sub>-GFP-Cdc3(R360A) | pRS306 | fl | This study |
| #43 | P<sub>MET</sub>-Cdc12-GFP | pRS306 | fl | This study |
| #44 | P<sub>MET</sub>-Cdc12(R263A)-GFP | pRS306 | fl | This study |
| #45 | P<sub>MET</sub>-Cdc11-GFP | pRS306 | fl | This study |
| #46 | P<sub>MET</sub>-Cdc11(R247A)-GFP | pRS306 | fl | This study |
| #47 | P<sub>MET</sub>-Shs1-GFP | pRS306 | fl | This study |
| #48 | P<sub>MET</sub>-Shs1(R288A)-GFP | pRS306 | fl | This study |

The septin complexes were purified by IMAC via the His_6_-tag and immediately afterwards subjected to analytical size exclusion chromatography (SEC). The elution profile of the wildtype SEPT7-SEPT9 heterodimer contained besides the major dimer peak a second peak corresponding to monomeric His_6_-SEPT9, likely resulting from unequal expression of the subunits from the respective plasmids (Fig. 6A). Mutation of Arg(βb) to Ala in any of the two subunits shifted the peak entirely towards the retention volume of monomeric His_6_-SEPT9. The SEPT7 homodimer eluted exclusively as dimer after IMAC-purification and the retention volume was entirely shifted to the SEPT7 monomer upon mutation of Arg(βb) to Ala (Fig.6B). Does the Arg(βb) to Ala mutation interfere with the process of G-interface formation? To answer this question, we assembled the SEPT7-SEPT9 heterodimer and the SEPT7 homodimer from individual subunits *in vitro* and tested if the Arg(βb) mutation inhibits complex formation. We generated His_6_-MBP-SEPT7 with and without Arg(βb) mutation and His_6_-SEPT9, His_6_-SEPT9(R(βb)A) and His_6_-SEPT7. All constructs were subsequently purified by IMAC and preparative SEC. The SEPT7 constructs tended to form dimers to a certain extent but addition of 1 mM MgCl_2_ to the IMAC elution buffer resulted almost exclusively in monomers as judged by SEC (Suppl. Fig. S9A).

**Figure 6.**
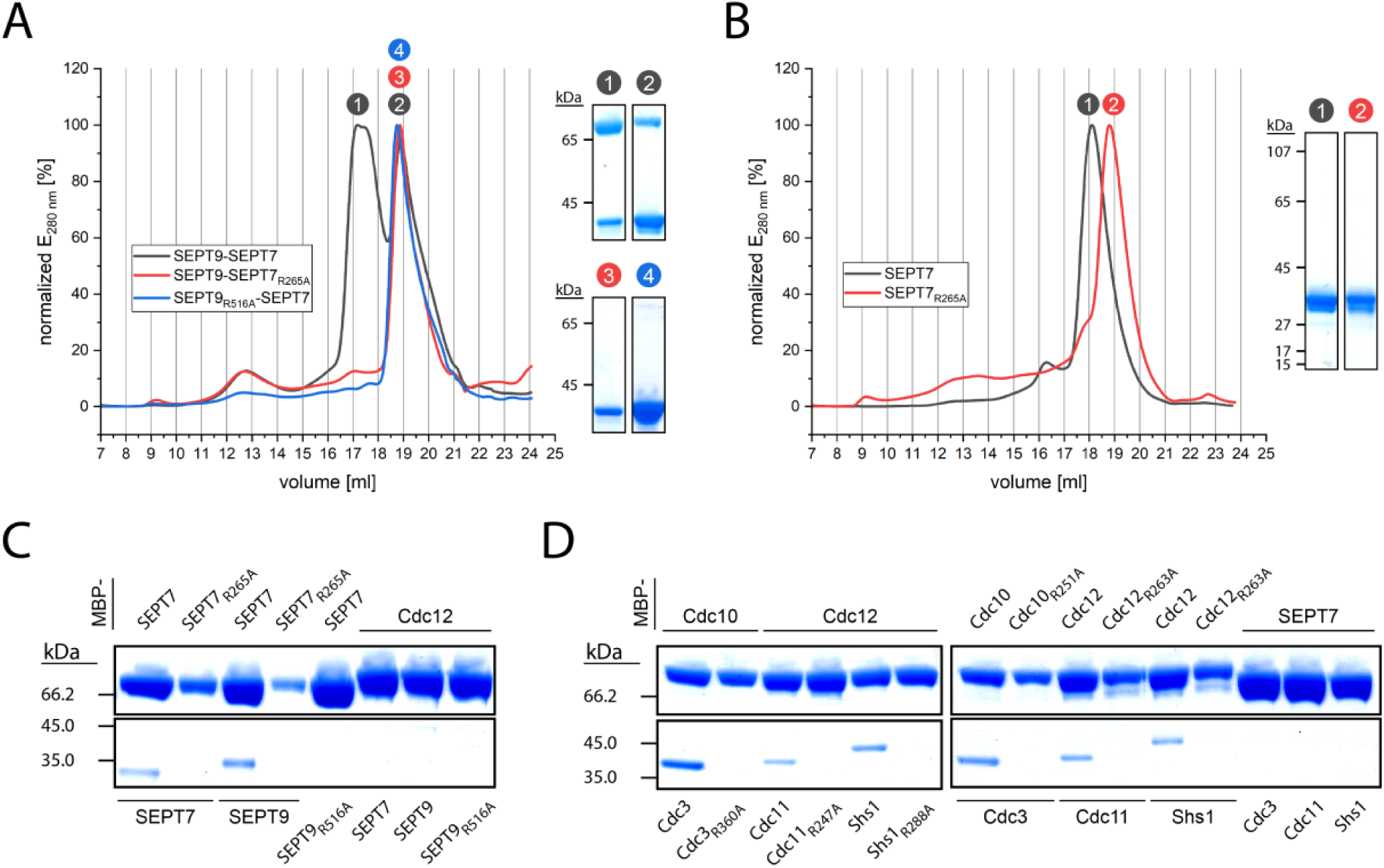
Arg(βb) is essential for nucleotide mediated G-interface formation *in vitro*. **(A)** His_6_-SEPT9 and MBP-SEPT7 G-domains with and without Arg(βb) mutation were co-expressed in E. coli, purified by IMAC and subsequently subjected to analytical SEC. Arg(βb) mutation in any of the subunits leads to disruption of the interface, leaving only monomeric His_6_-SEPT9 after the IMAC. Coomassie-stained SDS-PAGE sections of the indicated peaks from the SEC are shown besides the chromatogram. MBP-SEPT7: 75 kD calculated MW, His_6_-SEPT9: 36 kD. **(B)** His_6_-SEPT7 with and without Arg(βb) mutation was expressed in E. coli, purified by IMAC and subsequently subjected to analytical SEC. Arg(βb) mutation leads to disruption of the homotypic interface, leaving only monomeric His_6_-SEPT7 after the IMAC. Commassie-stained SDS-PAGE sections of the indicated peaks from the SEC are shown. His_6_-SEPT7: 33 kD calculated MW. **(C)** Purified monomeric MBP-SEPT7 with and without Arg(βb) mutation was incubated with purified monomeric His_6_-SEPT7, His_6_-SEPT9 and His_6_-SEPT9(R516A) in the presence of a GTP/GDP mixture. The Arg(βb) mutation suppressed G-interface formation entirely. Unspecific binding of human septin subunits to MBP-Cdc12 could not be detected. **(D)** Purified monomeric MBP-Cdc10 and MBP-Cdc12 with and without Arg(βb) mutation were incubated with the respective purified His_6_-tagged G-interface partner with or without Arg(βb) mutation in the presence of a GTP/GDP mixture. The Arg(βb) mutation suppressed G-interface formation entirely, regardless of the tested combination. Unspecific binding of yeast septin subunits to MBP-SEPT7 could not be detected.

The His_6_-MBP-SEPT7 (and His_6_-MBP-Cdc12) constructs were incubated with a five-fold molar excess of the respective His_6_-tagged G-interface partner in the presence of nucleotide.

Complexes were subsequently incubated with amylose resin, eluted with maltose and the composition of the complex was analyzed by SDS-PAGE. His_6_-SEPT9 and His_6_-SEPT7 were only able to bind to wildtype His_6_-MBP-SEPT7 (Fig. 6C). The Arg(βb) to Ala mutation suppressed G-interface formation entirely. No unspecific binding with the yeast septin His_6_-MBP-Cdc12 could be detected.

To test the G-interface formation for yeast septins, we purified monomeric His_6_-MBP-Cdc12 and His_6_-MBP-Cdc10 with and without Arg(βb) to Ala mutation and the respective G-interface partners His_6_-Cdc3, His_6_-Cdc11 and His_6_-Shs1. The monomeric state of the proteins was confirmed by SEC (Suppl. Fig S9B). While the G-domains of Cdc3, Cdc11 and Shs1 could be readily purified as monomers, Cdc10 and Cdc12 required the MBP-tag and an improved lysis buffer composition to ensure solubility (see Materials and Methods section).

As for the human G-interface dimers, the Arg(βb) to Ala mutation blocked formation of the Cdc10:3, Cdc12:11 and Cdc12:Shs1 G-interfaces entirely (Fig. 6D). No unspecific binding to the human His_6_-MBP-SEPT7 could be detected.

## Discussion

Small GTPases of the Ras family share common structural features and cycle between GTP and GDP bound conformations, each conformation exhibiting distinct affinities for effector molecules (Bourne *et al*., 1991). Although septins share the same structural features as small GTPases, their biochemistry seems to be different (Grupp and Gronemeyer, 2022) and the role of nucleotide utilization by septins is a subject of intense debate. The fact that all available dimeric or multimeric septin complexes contain nucleotide strongly suggests that septins utilize the nucleotide for protofilament formation (Grupp and Gronemeyer, 2022). We have previously employed MD simulations on available human septin structures to bypass the limitation that dimers in their apo conformation cannot be generated *in vitro* (Grupp *et al*., 2023). We found that an arginine residue in the SUE βββ-meander undergoes a conformational change upon nucleotide removal in the catalytically active SEPT2 and SEPT7 subunits. We could identify Arg(βb) related structural alterations also in yeast septins employing MD simulations.

When interpreting the output from MD simulations, one has to keep two intrinsic limitations in the context of septin dimers in mind: First, the simulations with septin dimers depict the scenario of disrupting an already formed dimer, but septin protofilaments are stable for hours *in vitro* once they are assembled (Renz *et al*., 2013). The simulations thus identify “weak spots” inside the dimer that also may determine dimer formation. However, these predictions have to be confirmed experimentally.

Second, the unbiased simulations cover a limited time frame (here 700 ns, requiring approx. two weeks of computing time per simulation) but folding of proteins or rearrangements within complexes may require microseconds (Chen *et al*., 2017), time scales exceeding reasonable computing time for unbiased MD. In our previous work, we argued that Arg(βb) flipping is a feature of catalytically active septins since we did not observe it in SEPT6. Contrary to this assumption, we could not show Arg(βb) flipping in the central clustered structure of the catalytically active Cdc12. Thus, we might have simply missed the flipping in Cdc12 as well as in the catalytically inactive Shs1 and SEPT6 if it occurs later on the reaction coordinate than what is captured by the time frame of the simulation.

To account for this presumable limitation, we have conducted free energy landscape and backbone RMSD analyses to detect and describe Arg(βb) related structural alterations in the SUE on a more global scale rather than to focus on the movement of one particular amino acid residue.

We performed *in vitro* and *in vivo* reconstitution experiments with human and yeast septins to confirm the evolutionary conserved significance of Arg(βb) for G-interface integrity.

SEPT9 is catalytically active and occupies the central position of the human octameric protofilament (Castro *et al*., 2020), making this subunit an excellent target to study Arg(βb) mediated filament incorporation of a subunit inside the living cell. We made use of our recently presented SEPT9 ko cell line (Hecht *et al*., 2024) and performed complementation experiments using EGFP-SEPT9 with and without mutated Arg(βb). The human septin cytoskeleton can consist of hexamers *in vivo* (Sellin *et al*., 2011; Soroor *et al*., 2020), with SEPT7 forming a central G-interface. Our SEPT9 ko cells show indeed a rudimentary septin cytoskeleton but display severe phenotypes such as defects in cell migration and focal adhesion- and filopodia dysfunction (Hecht *et al*., 2024). SEPT9 with Arg(βb) mutated to alanine is not incorporated into septin filaments *in vivo* and our *in vitro* data support the conclusion that the SEPT7-SEPT9 G-interface is not properly formed. Cells expressing this mutant as only copy of SEPT9 show a ko phenotype regarding filopodia architecture and migratory behavior.

The outer subunits of the yeast septin protofilament Cdc11 and Cdc12 are essential for the cell (Hartwell, 1971) and the Arg(βb) mutation in these subunits is lethal. Shs1 is a non-essential septin that can be replaced by Cdc11 (Iwase *et al*., 2007; Garcia *et al*., 2011) and consequently neither the ko of this subunit nor its Arg(βb)-mutation is lethal.

Neither Cdc11, nor Shs1, nor Cdc12 are incorporated into septin structures *in vivo* when bearing the Arg(βb) mutation, supported by the decreased inter-subunit affinity in the Shs1-Cdc12 dimer upon nucleotide removal and/or introduction of the mutant.

For the inner subunits of the yeast septin protofilament Cdc3 and Cdc10, introduction of the Arg(βb) mutation leads only to temperature sensitivity although both subunits are also essential (Hartwell, 1971). Localization to septin structures at the budneck is weakened for both subunits with Arg(βb) mutation, but still detectable. In line with these findings, the intra-subunit affinity in the Cdc3-Cdc10 interface is only decreased for the apo form but not for the interface bearing the Arg(βb) mutation. The Cdc3 and Cdc10 subunits including the Arg(βb) mutation might have a remaining sufficient (but weakened) overall affinity to form a G-interface *in vivo* (maybe promoted by other intracellular factors or conditions), resulting in the partial complementation of the corresponding gene deletions.

However, this remaining affinity is obviously not sufficient to form the interface *in vitro*. In Shs1, Cdc11 and Cdc12 the destabilizing effects prevail and the interface is neither formed *in vivo* nor *in vitro*.

Sequence analysis from 131 Opisthokonta and 123 Archaeplastida and Chromista septins shows the conservation degree of Arg(βb) across species and kingdoms (Fig. 7A). The corresponding sequence collection was retrieved from (Delic *et al*., 2024). All evaluated 254 septins contain Arg(βb) in their SUE-βββ and the other residues in the hydrogen network around Arg(βb) show a high conservation score (Fig. 7A). The conservation score was superimposed onto the structure of our yeast septin teramer (Fig. 7B). Closer inspection of the nucleotide binding pocket in Cdc3 as example shows that all residues of the hydrogen bond network coordinating the nucleotide with Arg(βb) in the center show the highest conservation score (Fig. 7C). Since we have confirmed nucleotide coordination by Arg(βb) also experimentally as crucial factor for G-interface formation in septins from yeast and man, we conclude that Arg(βb) is the central hub for nucleotide coordination in all septins. We suggest that during evolution septins have repurposed the nucleotide from signal transduction to a tool to induce and/or report protofilament completion.

**Figure 7.**
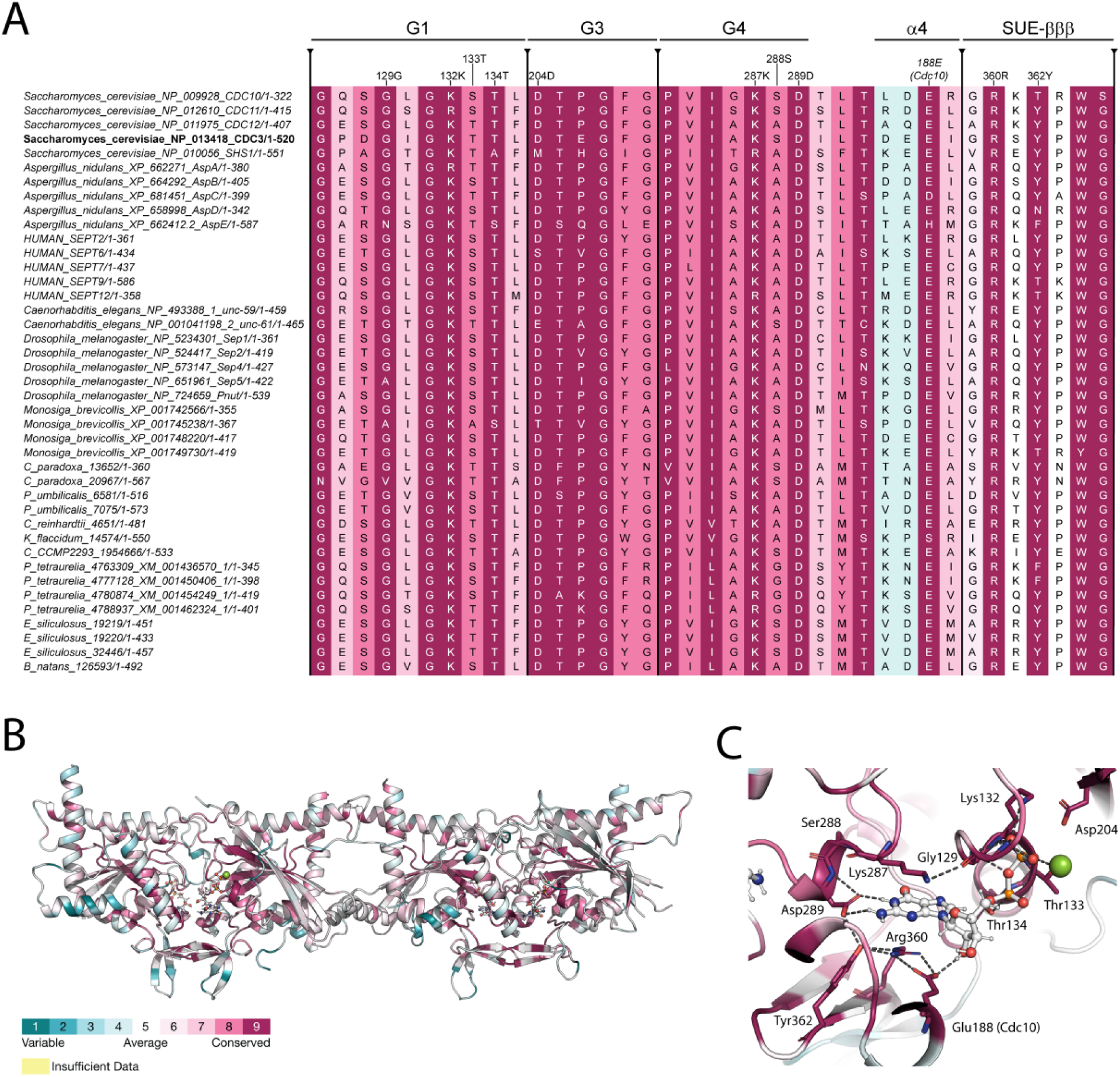
Sequence analysis from 254 septins (131 Opisthokonta and 123 non-Ophisthokonta). **(A)** Excerpt of the sequence alignment, showing 41 out of 254 septin sequences. Conservation degree is visualized by a color coded conservation score calculated for all 254 septins using ConSurf (Ashkenazy *et al*., 2010). Cdc3 (bold letters) serves as reference sequence and structure. **(B)** Conservation score from (A) projected on the yeast septin tetramer structure (PDB-ID 9GD4). The color gradient for the conservation score obtained from ConSurf is shown. **(C)** Nucleotide binding pocket of Cdc3. All amino acid residues within 5 Å distance from the nucleotide with conservation score of 9 are indicated. These residues are also indicated in (A) on the top of the alignment.

## Materials and Methods

### Protein crystallization and structure determination

Protein expression and purification for crystallization was performed as described previously (Grupp *et al*., 2024) with the exception that 10 mM NaF was added to the protein preparation after the preparative size exclusion chromatography.

Septin complexes were crystallized by sitting-drop vapor diffusion. Diffracting crystals were obtained with 0.33 µL of protein solution (3 mg / mL) mixed with 0.21 µL of a reservoir condition containing 17.5 % PEG 5000, 0.1 M (NH_4_)_2_SO_4_ and 0.1 M Bis-Tris pH 6.5 and 0.06 µL of a seeding solution. Crystals appeared within one day upon incubation of the crystallization plates at 20 °C. Crystals were cryoprotected in 10 % 2,3-butandiol prior to flash-freezing in liquid nitrogen. The diffraction experiments were carried out at the ID30B beamline of the ESRF (doi.10.15151/ESRF-ES-928402160) at 100 K at a wavelength of 0.873129 Å.

Merged X-ray diffraction data were obtained from the ESRF autoprocessing pipeline using XDS (Kabsch, 2010) as part of autoPROC Staraniso (Vonrhein *et al*., 2018).

The structure was solved by molecular replacement in Phaser (McCoy *et al*., 2007) using the subunits from PDB-ID 8PFH as template. The final model was completed by iterative rounds of automated refinement and model building in the Phenix package (Adams *et al*., 2010), autoBUSTER (Bricogne G., Blanc E., Brandl M., Flensburg C., Keller P. and Roversi P, Sharff A., Smart O.S., Vonrhein C., 2017) and Coot (Emsley *et al*., 2010). Initial automated refinement cycles in Phenix consisted of reciprocal and -real space positional refinement as well as grouped B-factor refinement and TLS refinement while applying secondary structure restraints. In later stages of model building automated refinement was performed using autoBUSTER with individual B-factor and TLS refinement. After refinement, the final model exhibited 97.85 % Ramachandran favored residues, of these 2.15 % in allowed regions, and a clash-score of 1.33 as determined by MolProbity as part of autoBUSTER.

The refinement statistics are shown as Suppl. Table 1.

### MD Simulations

Energy minimized coordinate files were generated as described in previous work (Grupp *et al*., 2023) with some modifications: The Cdc10:GDP-Cdc3:GDP:Mg^2+^ and Cdc12:GDP-Shs1:GDP dimers were extracted from PDB-ID 9GD4 and truncated to their G-domains (Cdc10_F31-I301_, Cdc3_F118-A410_, Cdc12_G33-E313_, Shs1_I22-S338_). Crystal waters within 0.6 nm of the truncated models were included for all further steps. Unresolved regions in the structures were filled using MODELLER (version 10.5) (Šali and Blundell, 1993) using subunits from PDB-ID 8SGD and 8PFH as templates. Assignment of protonation state, hydrogen bond optimization and structure relaxation were performed in Maestro (Schrödinger Release 2024-2, Maestro, Schrödinger, LLC, New York, 2021) as described previously (Grupp *et al*., 2023). Subsequently, the nucleotide coordinates were removed from the resulting PDB files, leading to structures of the dimers in the apo state. Both nucleotide-containing and apo coordinate files were generated with Arg(βb) to Ala mutations which were introduced with the “residue scanning” tool in Maestro.

Force field compatibility transformation as well as N- and C-terminal capping were performed essentially as described (Grupp *et al*., 2023).

All MD simulations were performed in GROMACS v. 2022.2 (Abraham *et al*., 2015) using the CHARMM36m force field (Huang *et al*., 2017) on the JUSTUS2 high performance cluster at Ulm University. Time steps for the integration were set to 2 fs and bonds to hydrogen atoms were constrained using the LINear Constraint Solver (LINCS) algorithm (Hess *et al*., 1997; Hess, 2008). No dispersion correction was applied. The particle mesh Ewald (PME) algorithm (Darden *et al*., 1998; Essmann *et al*., 1998) was employed for treatment of electrostatic interactions with a real-space cutoff of 1.2 nm. Short-range Lennard-Jones interactions were smoothly switched to zero between 1.0 and 1.2 nm. Periodic boundary conditions were applied in all directions.

Each dimeric complex was placed in an octahedral box with at least 1.5 nm distance between the protein and the surrounding box in all dimensions. The box was explicitly solvated using the CHARMM-modified TIP3P water model (Durell *et al*., 1994; Jorgensen *et al*., 1998; Neria *et al*., 1998). The system was further supplemented with 300 mM KCl, including neutralizing counterions. System construction was followed by steepest descent potential energy minimization of the system with a maximum force tolerance of 500 kJ mol^-1^ nm^-1^. Subsequently, equilibration to a target temperature of 300 K was performed over 50 ps under an NVT ensemble using the velocity rescaling thermostat (Bussi *et al*., 2007). The time constant for temperature regulation was set to 0.1 ps. Initial velocities were assigned using random seeding according to the Maxwell-Boltzmann distribution. Equilibration was continued for 500 ps under an NPT ensemble to reach the target pressure (1.0 bar). Pressure was regulated isotropically with the stochastic cell rescaling barostat (Bernetti and Bussi, 2020) with a time constant of 2 ps for pressure coupling. During temperature and pressure equilibration coordinates of heavy atoms in proteins and ligands were restrained with a force constant of 1,000 kJ mol^-1^ nm^-2^ in all dimensions. Production simulations were conducted under the same conditions as the NPT equilibration but without restraints and with pressure regulated by the Parrinello-Rahman barostat (Parrinello and Rahman, 1998). We performed unbiased MD simulations for 700 ns, during which we expected to be able to cover the Arg(βb) side chain motions and molecular rearrangements. Equilibration and production simulations were performed five times with different random seeds during velocity generation, resulting in sets of five independent production simulations per system.

Structural rearrangements within the Arg(βb) hydrogen bond network after unbiased simulations were analyzed by considering distances between key residues of this network, plotted as free energy landscapes. The residue identities were as follows: Asp(G4): Cdc10_D182_, Cdc3_D289_, Cdc12_D182_, Shs1_D220_; Glu(α4): Cdc10_E188_, Cdc3_D295_, Cdc12_D188_, Shs1_D226_; Arg(βb): Cdc10_R251_, Cdc3_R360_, Cdc12_R263_, Shs1_R288_. Free energies were obtained after extracting the pairwise distances between Arg(βb) and Asp(G4) as well as Arg(βb) and Glu(α4) from the five independent 700 ns simulations for each system. The reference atoms were Cζ in Arg(βb), Cγ in Asp(G4) and Cδ in Glu(α4). Distances from the five simulations were combined, followed by two-dimensional kernel density estimation in OriginPro v. 2023 using the bivariate kernel density estimator bandwidth method and the exact estimation formula. From the obtained densities, relative free energy differences (ΔG) were calculated between each observed state and the state exhibiting the highest probability density across corresponding apo and holo systems.

For quantification of global structural deformation in the SUE-βββ, the backbone RMSD for the corresponding amino acids (Cdc10_S237-E261_, Cdc3_S345-D370_, Cdc12_S248-E273_, Shs1_S277-D298_) was calculated after least square fitting of the investigated subunit on the equilibrated input structure. This was performed for each performed simulation independently. For the generation of distribution functions, the RMSD values obtained in the five independent 700 ns simulations for each system were combined separately for each SUE-βββ.

Input structures for COM-pulling simulations for both dimers were generated as described previously (Grupp *et al*., 2023) by pooling the last 100 ns of the five independent unbiased simulations and backbone RMSD-based clustering (Daura *et al*., 1998) after least-squares fitting. Clustering was performed with a cutoff of 0.2 nm except for the complex Cdc10_R251A_:GDP-Cdc3_R360A_:GDP:Mg. for which the cutoff was set to 0.25 nm as the lower cutoff yielded two dominant clusters of equivalent size. For each system the central structure in the largest cluster was subsequently used for further simulations. Surrounding waters and closely associated ions within 0.6 nm around the protein complexes were also retained for the following steps. Resulting models were placed in a rectangular box with x/y/z dimensions of 20/10/10 nm aligning the filament axis with the x-axis. Subsequent box solvation, energy minimization and equilibration were performed as described for the unbiased simulations. After equilibration, COM-pulling simulations were carried out in the absence of position restraints. Analysis of different combinations of pull rate and spring constant showed a similar behavior as described for human septin dimers (Grupp *et al*., 2023) resulting in the use of the same spring constant (500 kJ mol^-1^ nm^-2^) and a pulling rate (0.0075 nm ps^-1^). Using these parameters low structural deformation and reproducible maximum forces were obtained. COM-pulling simulations, umbrella sampling and calculation of free energy profiles were again performed as described in our previous work but at a temperature of 300 K and sampling COM distances until approximately 8.5 nm (Grupp *et al*., 2023).

### Plasmids and strains

All plasmids used and/or created in this study are listed in Table 2. All generated plasmids were verified by sequencing.

Plasmids for cell culture were created by modifying the EGFP-SEPT9 expression plasmid #1, kindly provided by Elias Spiliotis (Univ. of Virigina, School of Medicine, VA, USA). Point mutations were introduced via suitable primers, making use of an internal BamH1 restriction site located close to the respective codons, yielding plasmids #2 and #3, respectively.

For co-expression studies of human septins in E.coli, the G-domains of SEPT7 and SEPT9 were cloned into the in-house generated pET15b derivatives pES and pAc-MBP, allowing for simultaneous expression in *E.coli* due to compatible antibiotic resistance cassettes (Amp and Kan, respectively) and origins of replication (pBR322 and p15A, respectively). Inserts with point mutation were generated by SOE-PCR (SEPT7) using available plasmids as template (Fischer *et al*., 2022) or amplifying from #2 (SEPT9).

For expression of monomeric human- and yeast septins for pulldown experiments, the respective G-domains were PCR amplified from available plasmid templates and cloned as His_6_- or His_6_-MBP tagged constructs into the in-house generated pET15b derivatives pES and pAc or into the first MCS of pET-Duet1 (Novagen). Inserts with point mutation were generated by SOE-PCR.

For growth assays in yeast, the ORF of Cdc10, Cdc3, Cdc12, Cdc11 and Shs1 with and without R(βb)(A) mutation were cloned into the centromeric plasmid pRS313 (Sikorski and Hieter, 1989) enabling expression under the control of a P_MET17_ promoter yielding plasmids #29-#38. Inserts with point mutation were generated by SOE-PCR.

The ORF of Cdc10, Cdc3, Cdc12, Cdc11 and Shs1 were cloned into the centromeric plasmid pRS316 (Sikorski and Hieter, 1989) bearing an Ura3 prototrophy and a P_MET17_ promoter, yielding plasmids #24-#28 . These rescue plasmids were transformed into the yeast strain JD47 (Dohmen *et al*., 1995) by standard methods.

The wild type copy of the respective septin in the resulting strains was replaced by a NAT cassette as described elsewhere (Dünkler *et al*., 2021) and plasmids #29-#38 were subsequently transformed into the respective ko strains bearing the rescue plasmids.

The strains were grown in YPD medium over-night and serial dilutions were subsequently spotted on suitable selection media containing 5-FOA to drive out the rescue plasmid and on respective control media without FOA.

For microscopy, the ORF of Cdc10, Cdc3, Cdc12, Cdc11 and Shs1 (with and without R(βb)(A) mutation) were cloned das GFP fusion into the plasmid pRS306 (Sikorski and Hieter, 1989) under the control of a P_MET17_ promoter, yielding plasmids #39-#48. The plasmids were linearized with StuI and subsequently transformed into the haploid yeast strains JD47 and ULM53 (Grinhagens *et al*., 2020) bearing a Shs1-mCherry fusion (Schneider *et al*., 2013), resulting in homologous recombination into the native Ura3 locus.

### Protein expression and purification

The following human septin dimers, truncated to their G-domains, were employed for co-expression, with the respective compatible plasmids in the same cell: His_6_-SEPT9G (#6) + MBP-SEPT7G (#8), His_6_-SEPT9G(R516A) (#7) + MBP-SEPT7G (#8), His_6_-SEPT9G (#6) + MBP-SEPT7G(R265A) (#9), His_6_-SEPT7G (#10, SEPT7 homodimer), His_6_-SEPT7G (R265A) (#11).

The respective septin G-domain hetero- and homodimers were expressed in E. coli BL21DE3 LOBSTR in super broth medium at 18°C for 22pprox.. 20 h after induction with 1 mM IPTG. Cells were resuspended in IMAC A buffer (25 mM Tris pH8.0, 300 mM NaCl, 5 mM MgCl_2_, 15 % (v/v) glycerol), supplemented with 0.1% Tween20 and e-complete protease inhibitor cocktail (Roche), and lysed by lysozyme treatment and sonication. The resulting extract was submitted onto a 5 ml HisTrap Excel column (Cytiva) which was subsequently washed with 3 % IMAC buffer B (25 mM Tris pH8.0, 300 mM NaCl, 200 mM imidazole, 5 mM MgCl_2_, 15 % (v/v) glycerol) until a stable A_280_ baseline was obtained. The His_6_-tagged protein complexes were then eluted with 100 % IMAC buffer B.

The product peak was pooled and directly submitted to analytical size exclusion chromatography on a Superose6 column (Cytiva) with 25 mM Tris pH8.0, 300 mM NaCl, 5 mM MgCl_2_ as mobile phase. The dimerization state of the eluting proteins was judged according to a gel filtration calibration standard (BioRad).

Septin monomers for pulldown were expressed as described above.

#6, #7, #10, #11, #12, #13, #18-#23 expressing cells were resuspended in IMAC mono-A buffer (50 mM Tris pH8.0, 500 mM NaCl, 5 mM MgCl_2_), supplemented with 0.1% Tween20 and e-complete protease inhibitor cocktail (Roche), and lysed by lysozyme treatment and sonication. The resulting extract was submitted onto a 5 ml HisTrap Excel column (Cytiva) which was subsequently washed with IMAC mono-A including 3 % IMAC buffer mono-B (50 mM Tris pH8.0, 500 mM NaCl, 200 mM imidazole, 5 mM MgCl_2_) until a stable A_280_ baseline was obtained. The proteins were eluted with 100 % IMAC mono-B. For all SEPT7 constructs (#10, #11, #12, #13), IMAC mono-B was supplemented with 1M MgCl_2_ to dissociate SEP7 homodimers into monomers.

Eluates of #12 and #13 tended to precipitate under these conditions and were centrifuged for 5 min at 16.000 xg immediately before SEC.

Cells expressing His_6_-MBP-Cdc10 (#14 or #15) and His_6_-MBP-Cdc12 (#16 or #17) were lysed in IMAC buffer lowsol-A (50 mM Tris-HCl pH 8.0, 500 mM NaCl, 5 mM MgCl_2_, 50 mM L-Arg, 50 mM L-Glu, 15% Glycerol, 0.1% Tween20) supplemented with e-complete protease inhibitor cocktail (Roche) and lysed by lysozyme treatment and sonication. The resulting extract was submitted onto a 5 ml HisTrap Excel column (Cytiva) which was subsequently washed with IMAC lowsol-A including 3 % IMAC lowsol-B (50 mM Tris-HCl pH 8.0, 500 mM NaCl, 5 mM MgCl_2_, 50 mM L-Arg, 50 mM L-Glu, 15% Glycerol, 200 mM imidazole 0.1% Tween20) until a stable A_280_ baseline was obtained. The proteins were then eluted with 100 % IMAC lowsol-B. The product peaks of all septin monomers were pooled and subjected to preparative SEC on a Superdex200 column (Cytiva) with 25 mM Tris pH8.0, 300 mM NaCl as mobile phase.

All purified proteins were assayed by SDS-PAGE on a 4-12% BisTris gradient gel (Life Technologies) with subsequent Coomassie brilliant blue staining.

### Pulldown experiments

To enable complex formation, 2 µM MBP-tagged protein was incubated with 10 µM corresponding His_6_-tagged protein in pulldown buffer (25 mM Tris pH8.0, 300 mM NaCl, 5 mM MgCl_2_, 100 µM GTP, 100 µM GDP) in a total volume of 500 µl and incubated o/n at 4°C.

The complexes were then captured onto 30 µl amylose resin (NEB Biolabs) previously equilibrated with pulldown buffer, washed, and subsequently eluted with 80 µl MBP-elution buffer (25 mM Tris pH8.0, 300 mM NaCl, 5 mM MgCl_2_, 10mM maltose).

The eluates were subjected to SDS-PAGE on a 4-12% BisTris gradient gel (Life Technologies) and subsequently stained with Coomassie brilliant blue.

### Mammalian cell culture and microscopy

1306 fibroblasts (immortalized human dermal fibroblasts, female Puerto Rican donor, kind gift of Sebastian Iben, Dept. of Dermatology, Ulm University Hospital, Germany) and a SEPT9 knock out cell line in the same cellular background, generated previously in our laboratory (clone 2c3, (Hecht *et al*., 2024)), were used throughout this study. Cells were maintained at 37 °C and 5 % CO_2_ in a humidified incubator in DMEM (Gibco) supplemented with 10 % FBS (Capricorn). Plasmid transfection was performed using Lipofectamine 3000 (Invitrogen) according to the manufacturer’s instructions. Stable cell lines were selected by treatment 750 μg/ml geneticin G418 (Formedium) 48h post transfection. The antibiotic concentration was lowered to 250 μg/ml after approx. two weeks and the cells were further maintained under these conditions. Immunfluorescence, transwell migration assays and microscopy were performed essentially as described in previous work (Hecht *et al*., 2024). The following antibodies were used in this study: The endogenous actin cytoskeleton was stained using AlexaFluor555-Phalloidin (Invitrogen, #A34055). The endogenous septin cytoskeleton was stained using an anti-SEPT8 antibody (produced in rabbit, kind gift of Michael Krauss, Molecular Pharmacology and Cell Biology Group, Leibnitz Research Center for Molecular Pharmacology, Berlin, Germany) combined with an AlexaFluor647 conjugated anti-rabbit secondary antibody produced in goat (Invitrogen, #21244). 1306 fibroblasts contain SEPT8 as member from the 1b group instead of SEPT6 (Hecht *et al*., 2019). The anti-SEPT8 antibody stained septin cytoskeleton colocalized with DsRed-SEPT9 or GFP-SEPT2 decorated septin filaments in 1306 fibroblasts (Suppl. Fig. S10), proving specificity of the antibody in the employed cell line. Filopodia were stained with an anti-VASP antibody produced in rabbit (Invitrogen, #MA5-35728) combined with an AlexaFluor647 conjugated anti-rabbit secondary antibody produced in goat (Invitrogen, #21244).

For Western blot analysis, confluent cells from one T25 cell culture flask were collected and lysed in 100 µl lysis buffer (50 mM HEPES pH 7.4, 150 mM NaCl, 0.2 mM EDTA, 0.1 % Triton X100) supplemented with protease inhibitor cocktail (Roche). Cell debris was removed by centrifugation for 20 min at 16.000 xg at 4 °C and the protein concentration in the soluble extract was determined by Bradford assay. Equal amounts of protein for each tested cell lysate were loaded onto a 4-12% BisTris gradient gel (Life Technologies) which was subsequently blotted onto nitrocellulose. SEPT9 was detected using an anti-SEPT9 antibody (Sigma, #WH0010801M1) and an anti-GAPDH-HRP conjugate (Santa Cruz Biotechnology; #sc47724) was employed as loading control.

### Microscopy of yeast strains

Strains were grown over-night in suitable selection medium, diluted subsequently in 3 ml fresh medium, and grown for 1.5-2 h at 30 °C to mid-log phase. Directly before the microscopy analysis 1.5 ml of the cultures were spun down and resuspended in 100 μl sterile H_2_O. 3.5 μl of the suspension were spotted on a glass slide and immobilized with a coverslip. Microscopy was performed with a 63x magnification objective with a series of ten equally spaced z-slices, each separated by 0.26 μm. The same microsope setup as for the immunfluorescence was used (Hecht *et al*., 2024). Image analysis was conducted in Fiji (Schindelin *et al*., 2012). Sum projections of the acquired Z-planes from both channels were generated for subsequent analysis. To quantify bud neck signals, an elliptical region of interest (ROI) measuring exactly 8 pixels in length and 6 pixels in width was positioned over the bud neck, aligning the longer axis with the Shs1-mCherry signal and the shorter axis with the mother-daughter axis. Cytoplasmic signals in the mother and daughter cells were measured by placing an elliptical ROI over the remaining cell volume of the mother and daughter cells, respectively. Care was taken that these ROIs didn’t incorporate bud neck associated fluorescence signal in either channel. Only cells in G2/M phase displaying clearly discernible Shs1-mCherry signal at the bud neck were considered. Finally, the average intensity within each ROI was calculated for both channels. The ability of septin variants to be incorporated in septin filaments at the bud neck was quantified using the following equation:

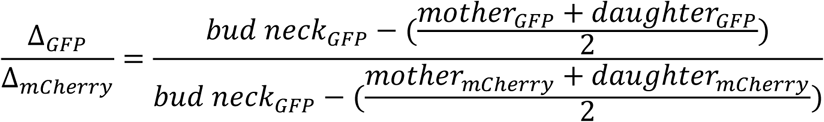

Using this equation, bud neck signals in both channels were corrected by subtracting the average cytoplasmic signals from the mother and daughter cells. GFP accumulation was then normalized relative to Shs1-mCherry accumulation.

## Data availability

Sequences of the employed oligonucleotide primers for cloning are available from the authors upon request. The refined structure of the yeast septin tetramer was deposited in the PDB under identifier 9GD4. Raw data underlying the figures including the full alignment containing all 254 sequences (Fig. 7), sequences of recombinant DNA and the MD data that support the findings of this study are openly available in Zenodo at doi. 10.5281/zenodo.14622194

## Acknowledgements

The authors thank Elias Spiliotis (Univ. of Virigina, School of Medicine, VA, USA), Matthias Hecht (formerly Institute of Molecular Genetics and Cell Biology, Ulm University) and Michael Krauss (Molecular Pharmacology and Cell Biology Group, Leibnitz Research Center for Molecular Pharmacology, Berlin, Germany) for the donation of plasmids and the anti-SEPT8 antibody, respectively.

The staff at beamline ID30B at the European Synchrotron ESRF is acknowledged for excellent assistance with data collection and automated processing.

The authors acknowledge support by the state of Baden-Württemberg through bwHPC, the German Research Foundation (DFG) through grant INST 40/575-1 FUGG (JUSTUS 2 high performance computing cluster) and the staff of the Uni Ulm computer center for technical support. Julia Seifermann is supported by DFG grants SFB1381 and RTG2202.

## Author Contributions

BG performed the simulations and experiments, built the structure, improved the manuscript and designed research. JBG performed experiments. JS crystallized the protein, collected and processed the diffraction data, and analyzed the data. SG analyzed the data.

JAL supported in setting up the simulations, analyzed the data and improved the manuscript. JFG performed experiments. NJ analyzed the data and improved the manuscript. TG analyzed the data, designed research, performed experiments and wrote the manuscript.

