## Supplementary information for "Interface integrity in septin protofilaments is maintained by an arginine residue conserved from yeast to man"

**Supplementary Table 1.** Crystallographic data collection and refinement statistics.

| | Cdc10-Cdc3-Cdc12-Shs1 |
|---|---|
| **Data collection** | |
| Space group | $P\ 1\ 2_1\ 1$ |
| Cell dimensions | |
| $a, b, c$ (Å) | 98.205, 47.845, 148.885 |
| $\alpha, \beta, \gamma$ (°) | 90.000, 103.286, 90.000 |
| Resolution (Å)[a] | 144.900 - 2.039 (2.355 - 2.039) |
| $R_{merge}$ | 0.304 (1.567) |
| $R_{meas}$ | 0.338 (1.753) |
| $R_{pim}$ | 0.145 (0.774) |
| $I / \sigma I$ | 5.2 (1.9) |
| Completeness spherical (%) | 48.2 (6.9) |
| Completeness ellipsoidal (%) | 89.3 (61.7) |
| Redundancy | 5.4 (5.0) |
| $CC_{1/2}$ | 0.973 (0.431) |
| **Refinement** | |
| Resolution (Å) | 23.49 - 2.039 (2.24 - 2.04) |
| No. reflections | 41834 (837) |
| $R_{work}$ / $R_{free}$[b] | 0.2161 / 0.2535 |
| No. of non-hydrogen atoms | |
| Protein | 9039 |
| Ligand/ion | 118 |
| Water | 586 |
| Mean $B$-factor (Å$^2$) | |
| Protein | 27.14 |
| Ligand/ion | 18.58 |
| Water | 23.37 |
| r.m.s. deviations from ideal values | |
| Bond lengths (Å) | 0.0082 |
| Bond angles (°) | 1.24 |
| Ramachandran statistics[c] | |
| Preferred regions (%) | 97.85 |
| Allowed regions (%) | 2.15 |
| Outliers (%) | 0.00 |
| Molprobity clash score, all atoms[c] | 1.33 |

A single crystal was used for the structure. Values in parentheses are for the highest-resolution shell.

[a]Data were truncated along the surface defined by $I/\sigma(I) = 1.2$ resulting in anisotropic diffraction limits of 2.645 Å, 3.018 Å and 2.017 Å along the reciprocal axes (0.745 a* − 0.667 c*), b* and (0.266 a* + 0.964 c*), respectively.

[b]Determined by MolProbity as part of autoBUSTER.

[c]Rfree was calculated as the Rwork for 5% of the reflections that were not included in the refinement.

A

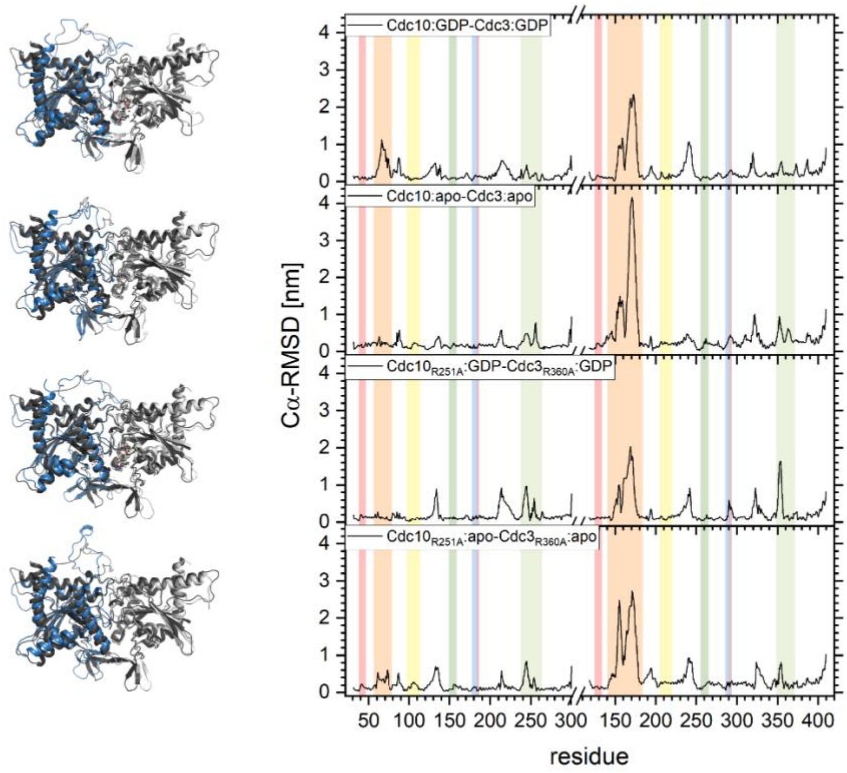

B

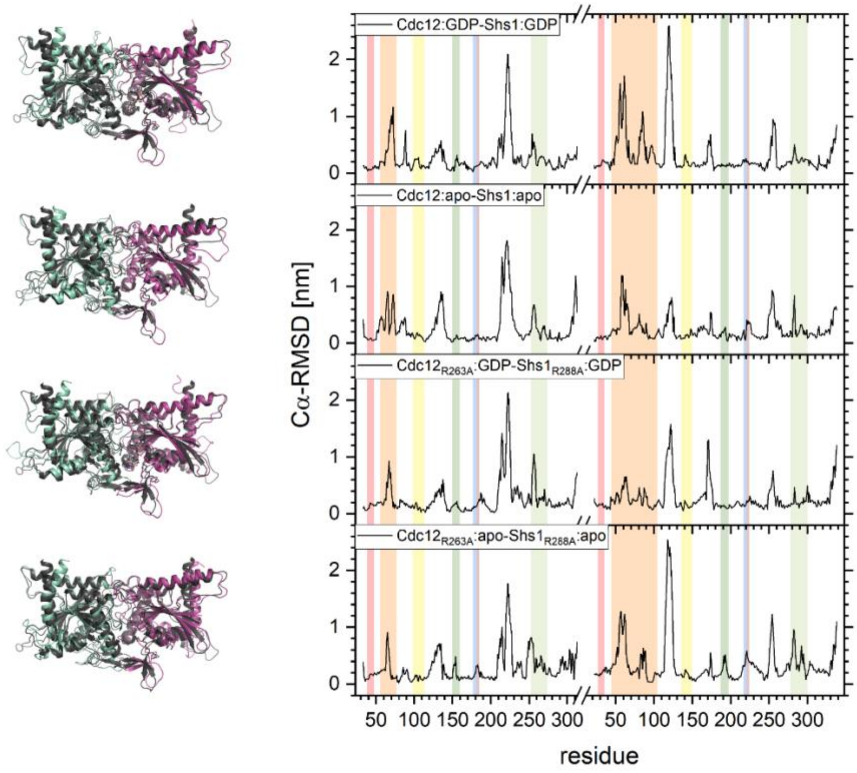

**Supplementary Figure S1**. Cα-RMSD of the relaxed structures compared to the energy minimized coordinate file structures for the Cdc10-Cdc3 **(A)** and Cdc12-Shs1 **(B)** G-interface dimers. For each system an overlay of the clustered, relaxed structure (blue/gray) with the coordinate file structure (dark-gray) (left) and a RMSD plot (right) is shown. The coloration of structural elements in the RMSD plots follows (Grupp et al. 2023).

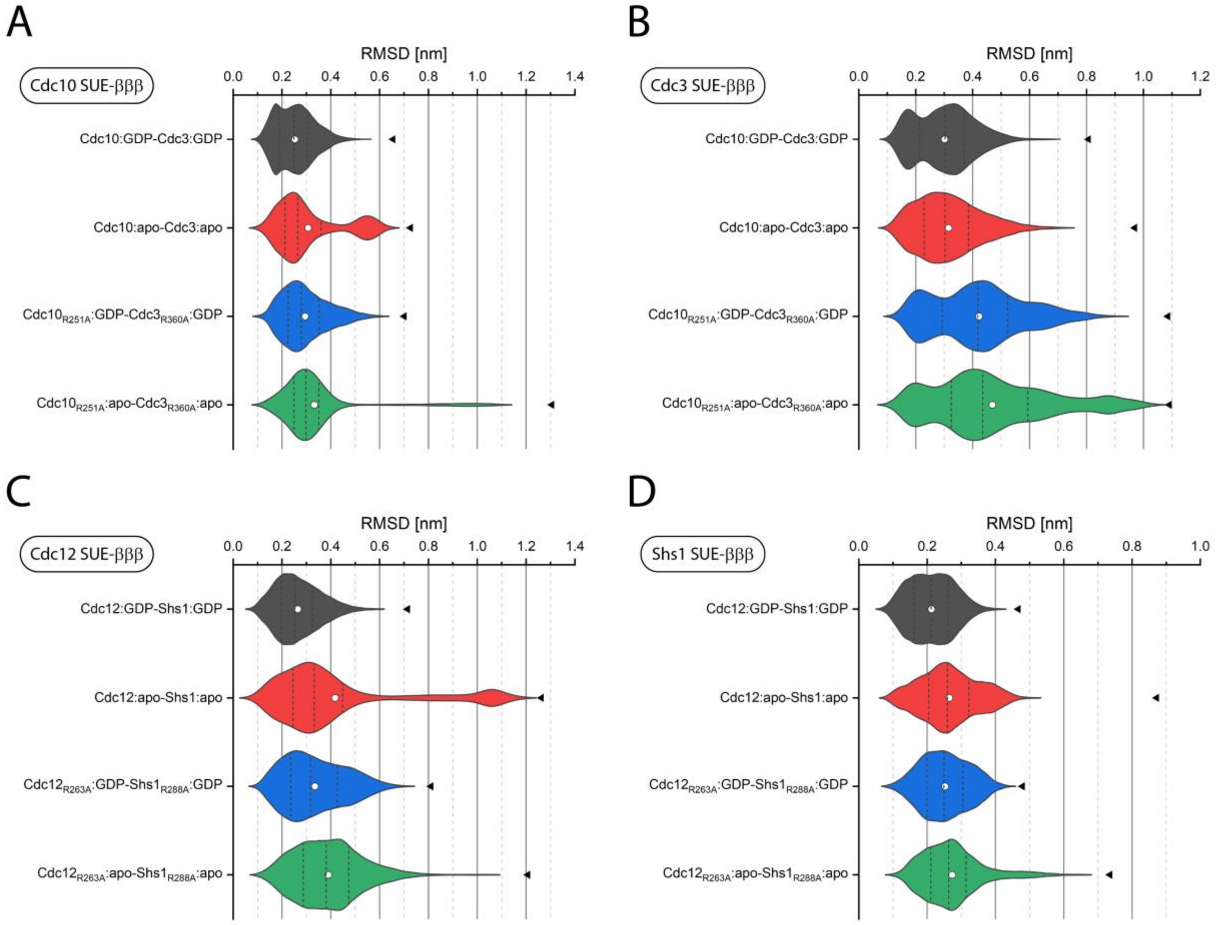

**Supplementary Figure S2**. Backbone RMSD analysis for the SUE-βββ. Shown is a RMSD distribution over the pooled unbiased simulations, with the input structures for the unbiased simulations (i.e. equilibrated coordinate structures) as reference. Black dashed lines represent the 25%, 50, 75% quantiles. The white dot indicates the average value of the RMSD distribution. The black arrowheads indicate the maximum RMSD value.
RMSD distributions are shown as violin plots for the SUE of Cdc10 (A), Cdc3 (B), Cdc12 (C) and Shs1 (D).

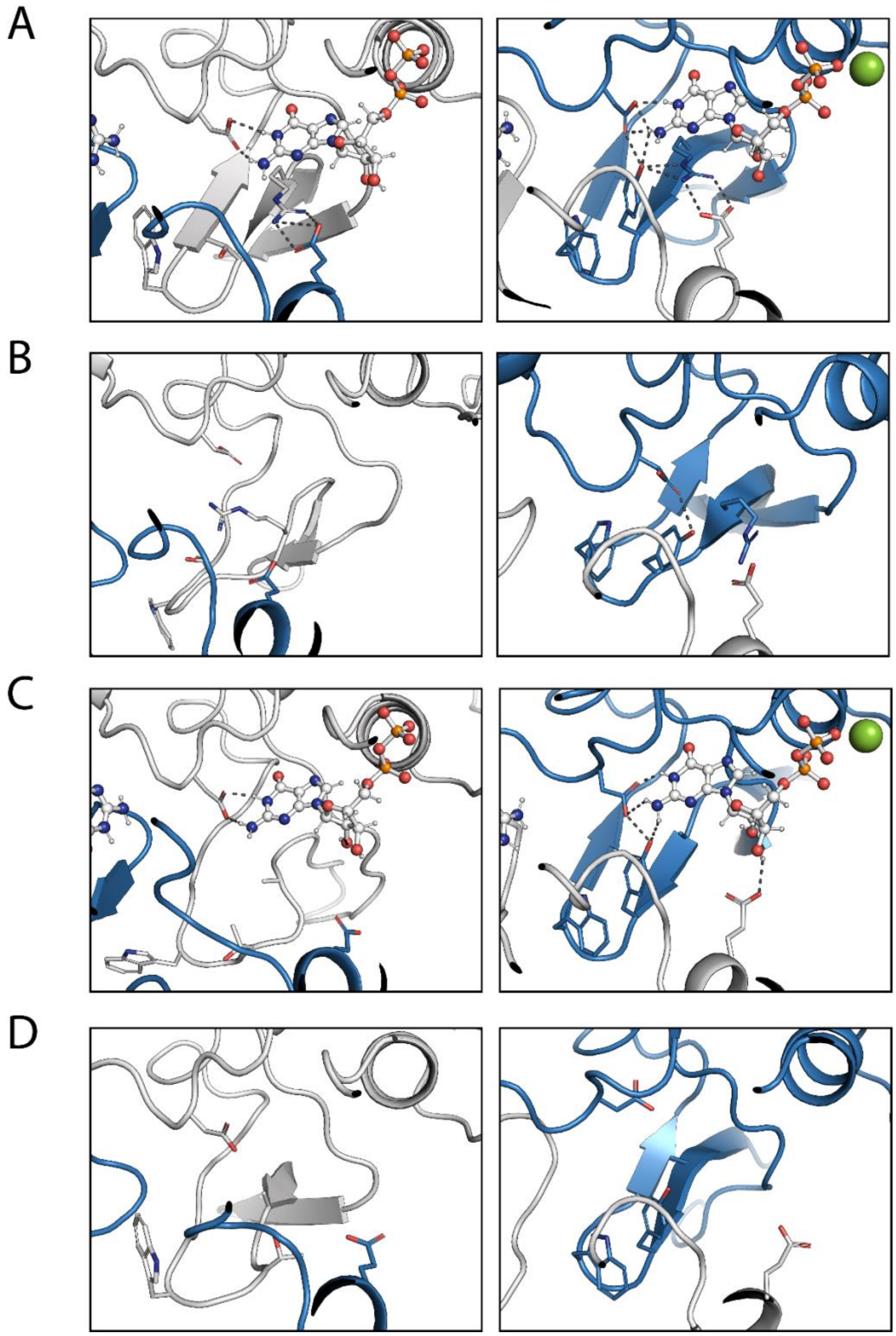

**Supplementary Figure S3.** Hydrogen bonding around the nucleotide in the different Cdc10 (gray) - Cdc3 (blue) systems, shown for the respective central structure of the largest cluster after unbiased simulations. **(A)** The hydrogen bonding network remains intact aft after unbiased simulations. **(B)** In the apo form, the hydrogen bond network becomes disturbed, going along with Arg(βb) flipping. **(C)** Upon Arg(βb)-Ala mutation, the hydrogen network in Cdc10-SUE(βbβc) becomes disturbed and the Cdc10-SUE(βββ) is partly dissociated from Cdc3. **(D)** The Arg(βb)-Ala mutation in the apo form leads also to perturbation of the Cdc10-SUE(βbβc) hydrogen bond network. The overall topology of the SUE remains intact.

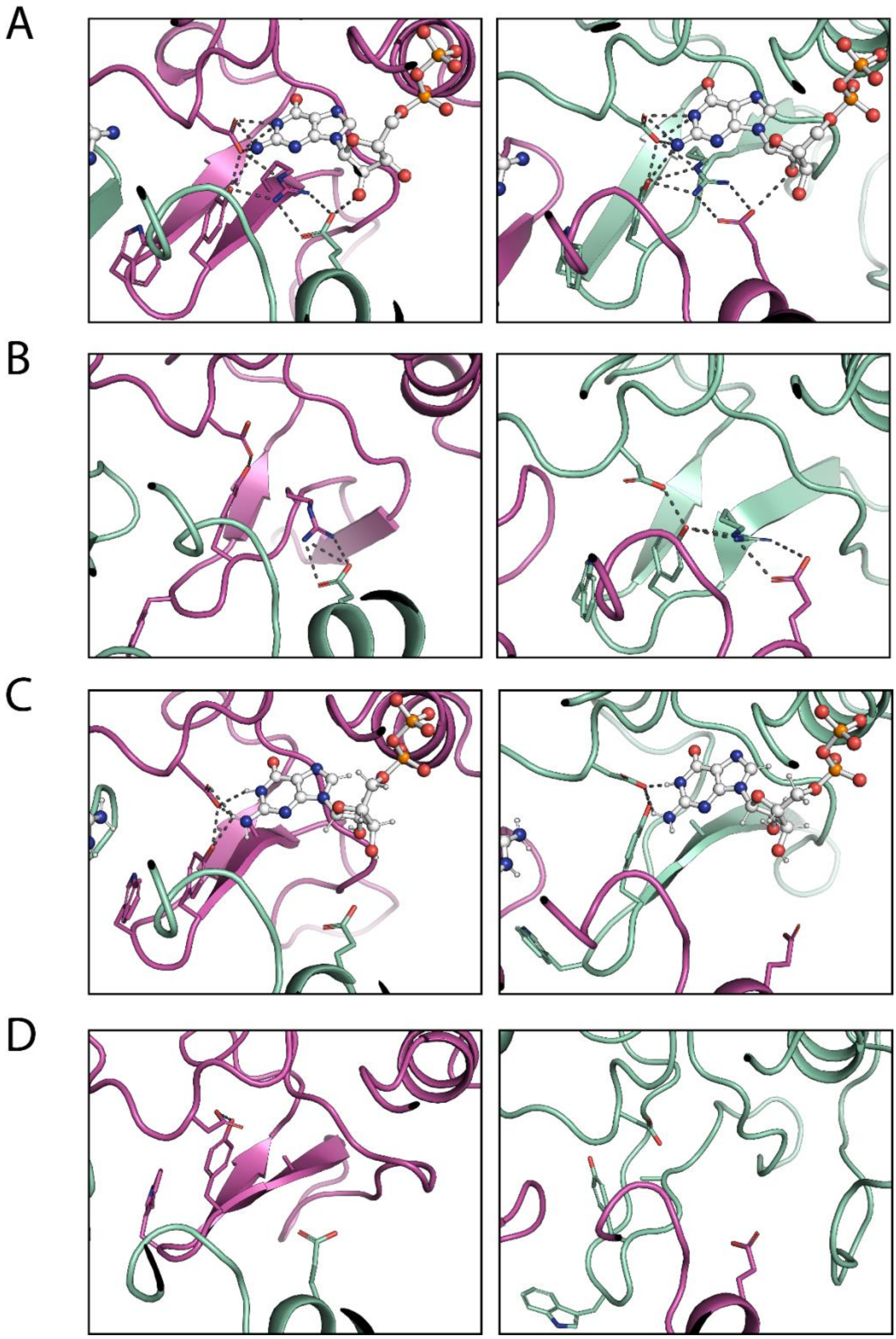

**Supplementary Figure S4.** Hydrogen bonding around the nucleotide in the different Cdc12 (pink) - Shs1 (pale-green) systems, shown for the respective central structure of the largest cluster after unbiased simulations. **A)** The hydrogen bonding network remains intact aft after unbiased simulations. **(B)** In the apo form, the hydrogen bond network around Arg(βb) becomes disturbed but no flipping can be seen. Tyr(βb) from Cdc12 tilts towards Trp(βb) from Shs1. **(C)** Upon Arg(βb)-Ala mutation, the hydrogen bond network becomes disturbed. **(D)** The Arg(βb)-Ala mutation in the apo form leads to partial dissociation of the Shs1-SUE(βββ) from Cdc12.

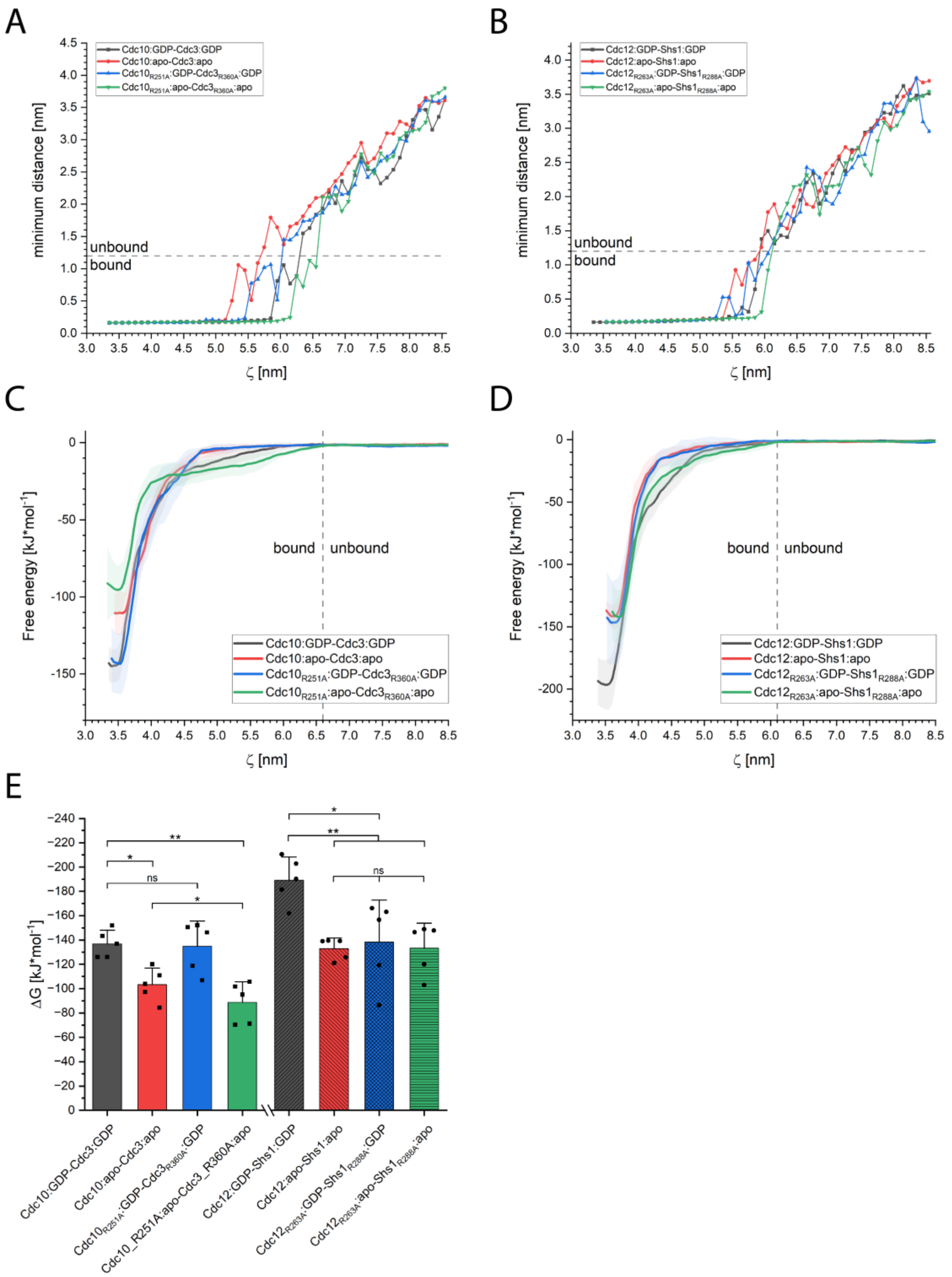

**Supplementary Figure S5.** Determination of binding free energies. **(A)** Median minimum inter-subunit distances along the reaction coordinates for the Cdc10-Cdc3 (left) and Cdc12-Shs1 systems (right), extracted from umbrella sampling windows. Dashed lines indicate the nonbonded interaction cutoff. ΔG was calculated from the Cα-COM distances separating the bound and unbound sections. **(B)** Average free energy profiles for the Cdc10-Cdc3 (left) and Cdc12-Shs1 systems (right). **(E)** Calculated ΔG values for all systems visualized as bar plot. Error bars represent the standard deviation. p-Values were determined via ANOVA followed by a Tukey post hoc analysis. Significant p-values are as follows: * p ≤ 5%, ** p ≤ 1%, *** p ≤ .1%, **** p ≤ .01%.

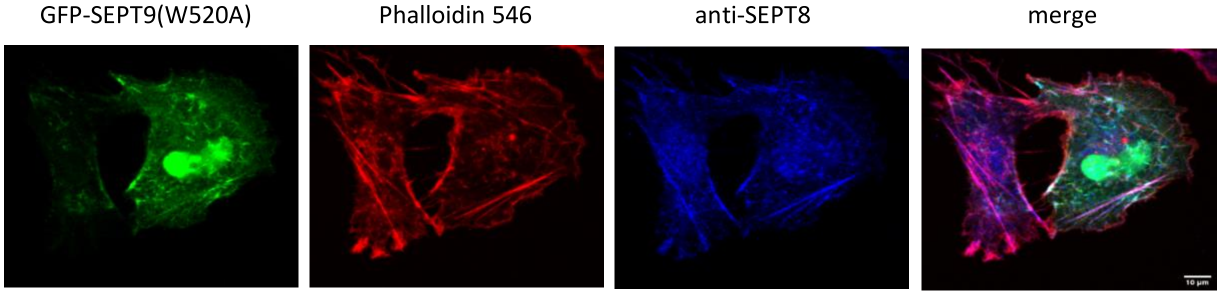

**Supplementary Figure S6.** Stable overexpression of GFP-SEPT9(W520A) in SEPT9 ko fibroblasts. The actin cytoskeleton was stained with Phalloidin and the endogenous septin cytoskeleton was stained with an anti-SEPT8a antibody (rabbit), followed by an anti-rabbit Alexa647 conjugated secondary antibody. Scale bar 10 µM.

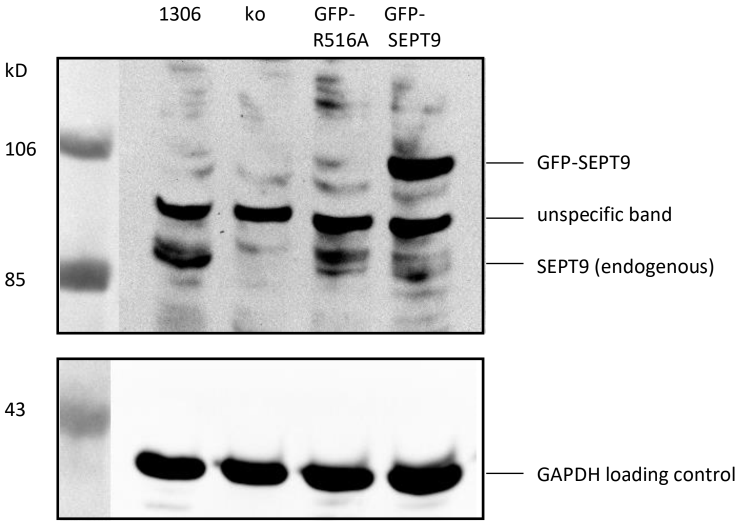

**Supplementary Fig. S7.** Expression analysis using an anti-SEPT9 antibody in soluble extracts from untreated 1306 fibroblasts, 1306 SEPT9 ko fibroblasts and SEPT9 ko fibroblasts expressing GFP-SEPT9(R516A) and GFP-SEPT9. Note a certain degree of degradation for the GFP-SEPT9 fusion proteins.

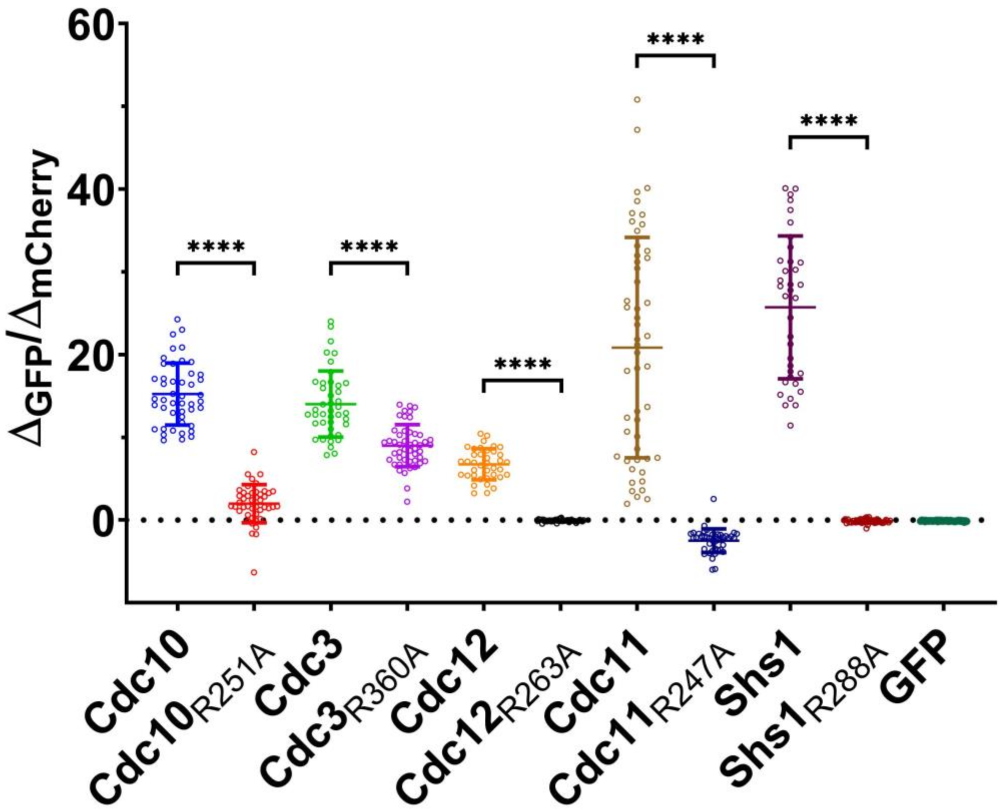

**Supplementary Figure S8**. Localization of septins containing Arg(βb) mutation inside the yeast strain ULM53. Septins with and without Arg(βb)-Ala mutation were expressed as GFP fusion protein in yeast cells containing Shs1-mCherry as septin marker. Shown is the quantification of the bud neck signal (in relation to the marker signal) of the GFP fusion constructs. Only cells in G2/M phase with a clearly detectable Shs1-mCherry signal at the bud neck were considered. Bud neck localization is entirely abolished for Arg(βb) mutation in Cdc12, Cdc11 and Shs1 and reduced for Cdc10 and Cdc3. Significance values were calculated by an unpaired t-test with Welch's correction with at least N=36 cells.

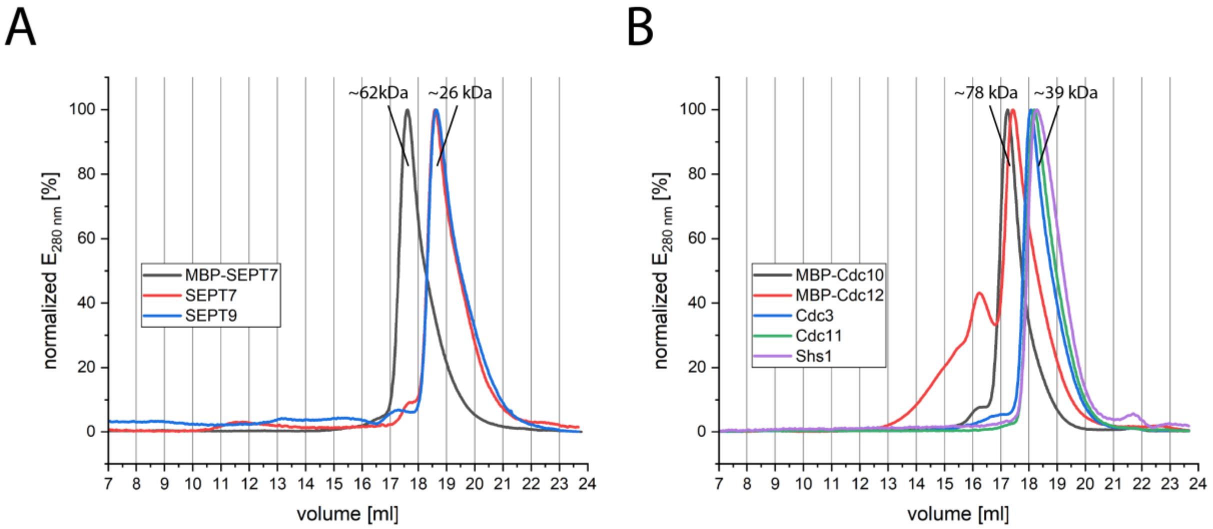

**Supplementary Figure S9**. Analytical SEC to evaluate the monomeric state of the septin subunits employed for pulldown experiments. The indicated molecular weight of the peaks is derived from a gel filtration standard containing five globular proteins. **(A)** G-domains of human SEPT7, SEPT9 and MBP-SEPT7 elute as monomers from analytical SEC. **(B)** G-domains of yeast Cdc3, Cdc11, Shs1, MBP-Cdc10 and MBP-Cdc12 elute as monomers from analytical SEC. MBP-Cdc12 tends to form auto-aggregates to a minor extent, indicated by the shoulder in the chromatogram.

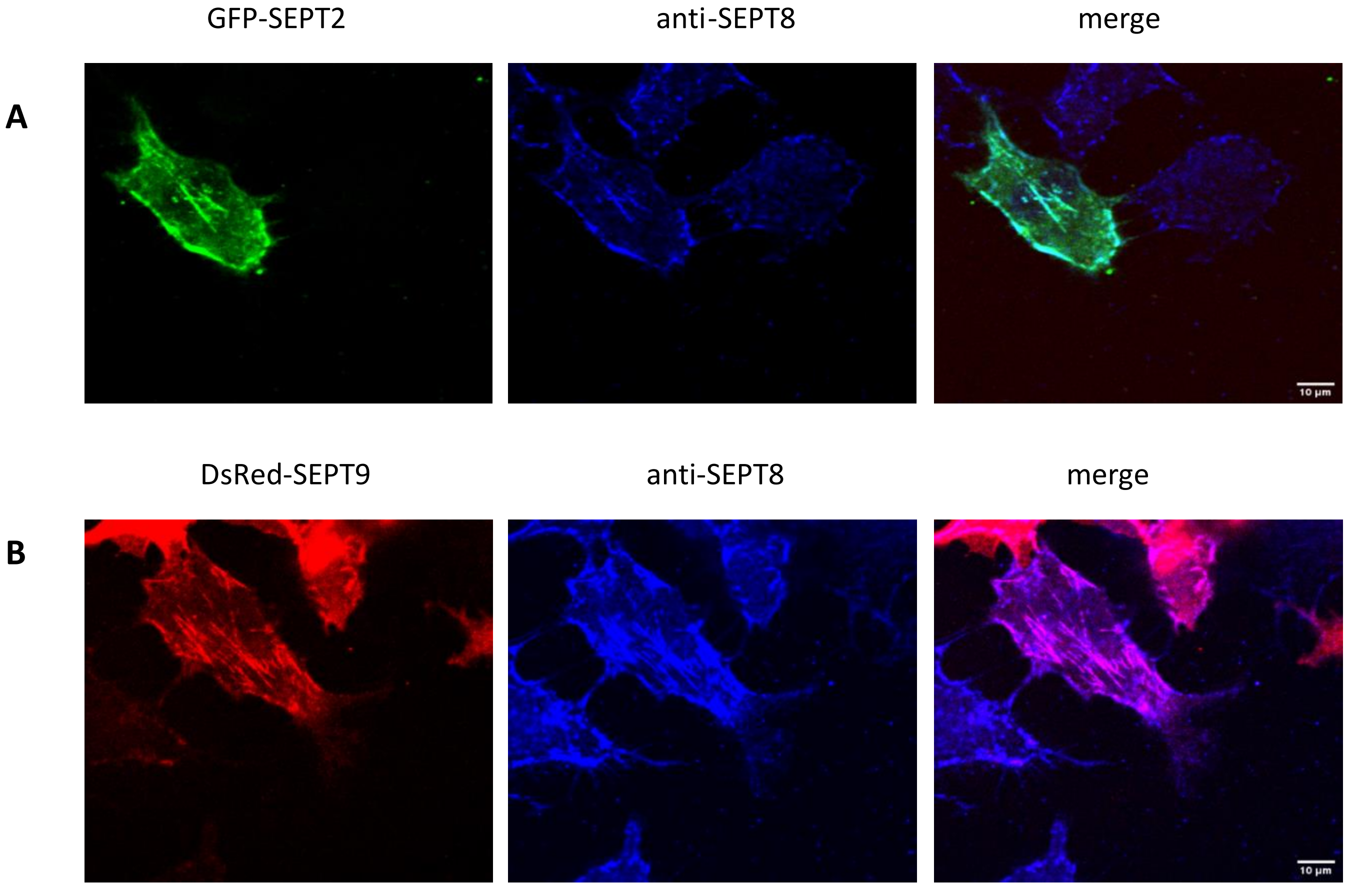

**Supplementary Figure S10.** Validation of the anti-SEPT8 antibody. 1306 fibroblasts were transiently transfected with **(A)** GFP-SEPT2 or **(B)** with DsRed-SEPT9 expression plasmids, respectively. SEPT8 was stained using the anti-SEPT8 antibody (produced in rabbit), combined with an anti-rabbit Alexa647 conjugate. Note the colocalization of both fusion proteins with SEPT8. Scale bar 10 µM.
